# Higher-order interaction inhibits bacterial invasion of a phototroph-predator microbial community

**DOI:** 10.1101/564260

**Authors:** Harry Mickalide, Seppe Kuehn

## Abstract

In nature, the composition of an ecosystem is thought to be important for determining its resistance to invasion by new species. Studies of invasions in natural ecosystems, from plant to microbial communities, have found that more diverse communities are more resistant to invasion. It is thought that more diverse communities resist invasion by more completely consuming the resources necessary for would-be invaders. Here we show that *Escherichia coli* can successfully invade cultures of the alga *Chlamydomonas reinhardtii* (phototroph) or the ciliate *Tetrahymena thermophila* (predator), but cannot invade a community where both are present. The invasion resistance of the algae-ciliate community is due to a higher-order (3-way) interaction that is unrelated to resource consumption. We show that the mechanism of this interaction is the algal inhibition of bacterial aggregation which leaves bacteria vulnerable to ciliate predation. This mechanism of invasion resistance requires both the algae and the ciliate to be present and provides an example of invasion resistance through a trait-mediated higher-order interaction.

---

A hypothesis in microbial ecology states that “everything is everywhere, but the environment selects.”^1^ Under this hypothesis most microbial species are present in most environments, and it is the particular qualities of a local environment that determine which species establish themselves. This hypothesis is supported by studies in oceans^2^ and streams^3^ which show that the composition of microbial communities is better predicted by local environmental variables than spatial location^4^.

The observation that local environmental variables strongly predict community composition suggest that rates of dispersal are high, and indeed from active transport like motility^5^ to passive transport like wind^6^ and water currents, there are many mechanisms by which microbes disperse. Moreover, rapid dispersal suggests persistent immigration of microbes into new communities. Whether or not a given species successfully invades and establishes itself when encountering a new environment depends not just on the environment, but also on the invasion resistance of the local microbial community.

Therefore, understanding the mechanisms of invasion resistance in microbial communities is central to understanding the structure of microbial communities in nature. Moreover, a fundamental understanding of microbial community invasion dynamics is necessary for successfully designing industrial processes such as algal biofuel production^7^ or controlling harmful invasions in nature such as microbial blooms^8^.

Our current understanding of when and why some invasions succeed and others fail is grounded in the idea that an invader must either outcompete an existing community member for an available resource (dominance) or consume a resource that is not already being consumed by the community (complementarity)^9^. Dominance and complementarity have successfully explained the outcome of invasions in a wide range of studies. For example, in laboratory populations of *Pseudomonas fluorescens*, more diverse communities resisted invasion more effectively by more completely occupying the available niches^10^. Qualitatively similar results were observed for *E. coli* invasions of soil communities^11^, wherein diversity of and resource consumption by the community were positively correlated with invasion resistance. Similarly, studies of plant root bacterial communities demonstrated that community resource competition networks could reliably predict the outcome of invasions both *in vitro* and in tomato plant root communities^12^. Similar results have been found for plant communities on a larger spatial scale^13^.

Recently, theoretical work using consumer-resource models has extended this intuition and suggested that the emergent resource consumption and exchange in cross-feeding communities can be understood as a community-level fitness which provides cohesiveness and therefore invasion resistance^14^. Experimental efforts suggest that this picture can capture some features of experimental invasions in bacterial communities^15^. Collectively, this work shows that substantial insight into invasion dynamics can come from understanding resource dynamics during an invasion process.

However, in nearly all microbial communities there exist interactions that are not directly mediated by resources: for example, antagonistic interactions such as the excretion of antibiotics^16^ and predation by protists^17^ or phage^18^. A handful of studies have examined the role these interactions play in determining the fate of invading species^19^, but, as recently pointed out by Mallon *et al.*^20^, it remains an outstanding question how antagonistic interactions affect community invasion dynamics.

Here we use a model microbial community to study invasion dynamics in the presence of antagonistic interactions. Microbial communities in freshwater lakes and nearby saturated soils are occupied by primary producers who fix inorganic carbon, metabolically flexible heterotrophic bacteria who decompose organic matter, and predators who unlock nutrients held in biomass^21^. To study this canonical natural community, we use a three-species model microbial ecosystem comprised of the alga *Chlamydomonas reinhardtii* which acts as a primary producer and is an endemic phototroph in soils and freshwater^22^, the bacterium *Escherichia coli* which acts as a decomposer and is common in soils^23^, and the ciliate *Tetrahymena thermophila* which dwells in freshwater and preys on *E. coli*. We refer to this model ecosystem as the ‘ABC’ community for **A**lgae, **B**acteria and **C**iliates. The ABC community has been studied previously as a model self-sustaining closed microbial ecosystem^24–26^. Recent work has shown that long-term abundance dynamics in closed ABC ecosystems are complex and deterministic on timescales of months, exhibiting rich spatiotemporal and phenotypic dynamics^25^. The fact that the composition of the ABC community reflects the structure of some natural communities and that quantitative measurements are feasible make this a compelling model ecosystem for quantitative ecology^27^.

Here we show that when *E. coli* (B) is introduced into communities of *C. reinhardtii* (A) and *T. thermophila* (C), a higher order (3-way) interaction determines the outcome of the invasion. When B invades C alone, B aggregates to avoid predation by C and successfully grows to high density. Similarly, when B invades A alone, A may stall the invasion of B, but B can still successfully invade and grow to high density. In contrast, when B is introduced into a community of C and high-density A (> 5 × 10^4^ mL^−1^), B always fails to invade. We demonstrate that nutrient competition is not responsible for the invasion dynamics we observe. Instead, we find that A inhibits aggregation of B, resulting in increased predation pressure on B by C and therefore a decline in B abundances.

## Results

We study the dynamics of the ABC community in batch culture conditions with the community open to gas exchange. Organisms are introduced at low initial densities into 30mL of a freshwater mimic medium^28^ with undefined carbon and nitrogen sources (proteose peptone No. 3, see Methods). To initiate an experiment, all three organisms are cultured axenically in their respective growth media. Cells are washed and then their densities determined by flow cytometry. The communities are then constructed with known initial starting densities and maintained in custom culture devices which control temperature via feedback to a Peltier element (30 °C) and illumination via an LED below the vial (Fig. 1a).

**Figure 1:**
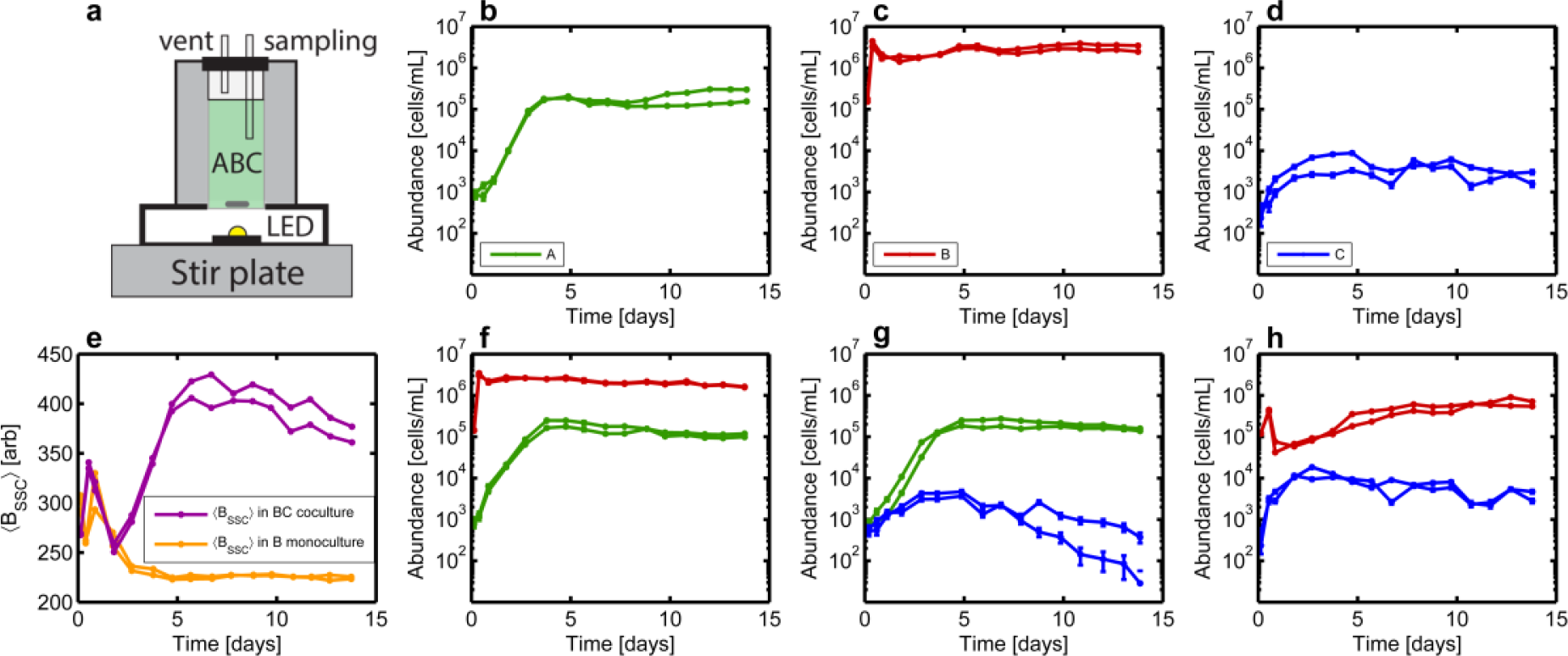
Monoculture and pair-culture dynamics with algae, bacteria, and ciliates. **a,** A schematic of the custom culture devices used in this study. **b-d,** Abundance dynamics plotted for monocultures of algae (A), bacteria (B), and ciliates (C) respectively at 4200 Lux (high light). **f-h,** Abundance dynamics for AB, AC, and BC pair-culture respectively, also at 4200 Lux. For each experiment there are two independent replicates. Abundances are measured via flow cytometry. Error bars are computed as described in Methods. For time points where error bars are not visible, errors are smaller than the size of the points. B abundances are reported as the total number of cells including planktonic cells and cells in aggregates as described in the SI. **e**, Mean side-scatter signal of B as a function of time in B monoculture and BC pair-culture.

Here we present two types of experiments: co-culture experiments and invasion experiments. Co-culture refers to experiments in which all species are simultaneously introduced at low densities (1 × 10^3^ mL^−1^ for A, 5 × 10^2^ mL^−1^ for B, and 5 × 10^2^ mL^−1^ for C) and in all possible monoculture, pair-culture, and tri-culture combinations. Invasion refers to experiments in which A and/or C is introduced at low density, allowed to grow for a fixed period of time (4 or 14 days), and then inoculated with B. In all experiments, abundance dynamics are followed approximately daily by sampling 500 µL of the community and performing flow cytometry measurements. Flow cytometry permits the quantification of abundances for A, B and C by chlorophyll fluorescence, genetically encoded yellow-fluorescent protein (YFP) fluorescence and size respectively (see Methods). Since a significant number of B cells are present in aggregates, we apply an aggregate correction algorithm (see SI) to estimate the true abundance of B cells and that is what we report in this study. We varied A’s growth rate by performing experiments at two light levels: ‘low light’ (average intensity of 1600 Lux) or ‘high light’ (average intensity of 4200 Lux). Abundance dynamics of B and C are not altered by illumination over this range of intensities (Fig. S1c&d, Fig. S8a). Communities are mixed by a magnetic stirrer at a rate of 450 rpm and sampled through a sterile port.

### Monoculture and pair-culture dynamics

To begin, we measured monoculture and pair-culture dynamics in the high light condition (Fig. 1). pair-culture dynamics between A and B suggest limited impact of A on B growth rate or carrying capacity (Fig. 1b,c,f). Similarly, B does not measureably impact A growth rate or carrying capacity in this high light condition (Fig. S1b).

In contrast, when B is pair-cultured with C, we observe an approximately 10-fold reduction in the abundances of B relative to B monoculture. This reduction is expected due to the known predation of B by C. In these BC pair-cultures, predation of B by C fails to drive B abundances below approximately 10^5^ mL^−1^ and at longer times B abundances increase (Fig. 1h). Previous measurements of ciliate feeding rates^29^, however, suggest that at these densities, C should be able consume most of the B present (see SI). We propose that the ability of B to sustain comparatively high densities in the presence of C is driven by B aggregation^30^. Bacterial aggregation is a common defense against predation due to the fact that the oral apparatus of the ciliates has a limited range of prey sizes it can accommodate ^31, 32^. Sufficiently large aggregates of B cannot be consumed by C. Indeed, we show that B aggregates much more in the presence of C than in monoculture (Fig. 1e). B aggregation was quantified by side-scatter measurements (Fig. S2). We conclude that B abundances are reduced by predation but the impact of predation is limited by aggregation. We also note that C abundances are not substantially impacted by the presence of B. This fact is in accordance with the low yield of ciliates on bacteria since previous work suggests 1 × 10^3^ to 4 × 10^4^ bacteria are required to produce a single ciliate (see SI).

Finally, when A and C are pair-cultured, the dynamics of C are minimally impacted relative to C monoculture (Fig. 1d,g). Taken together, Fig. 1 suggests that the dominant interaction in the ABC community is predation of B by C while interactions between A and B or A and C are limited.

### Bacterial invasions of ciliates

We studied the dynamics of B invading C. We introduced B at a density of ~ 1 × 10^4^ mL^−1^ into established cultures of C 4 and 14 days after the initiation of C cultures (Fig. 2a,d,g). We find that irrespective of the timing of the introduction of B or the light levels, B successfully grows to high densities (> 1 × 10^5^ mL^−1^). As in BC pair-culture (Fig. 1e), side-scatter intensity for B in invasion experiments confirms B aggregation in the presence of C (Fig. S6). For the purposes of discussion, we define a successful B invasion as one in which B abundances exceed 7 × 10^4^ mL^−1^ at the end of the experiment, or, in the case of panel **c,** when B abundances rise above 7 × 10^4^ mL^−1^ and then remain high for several days.

**Figure 2:**
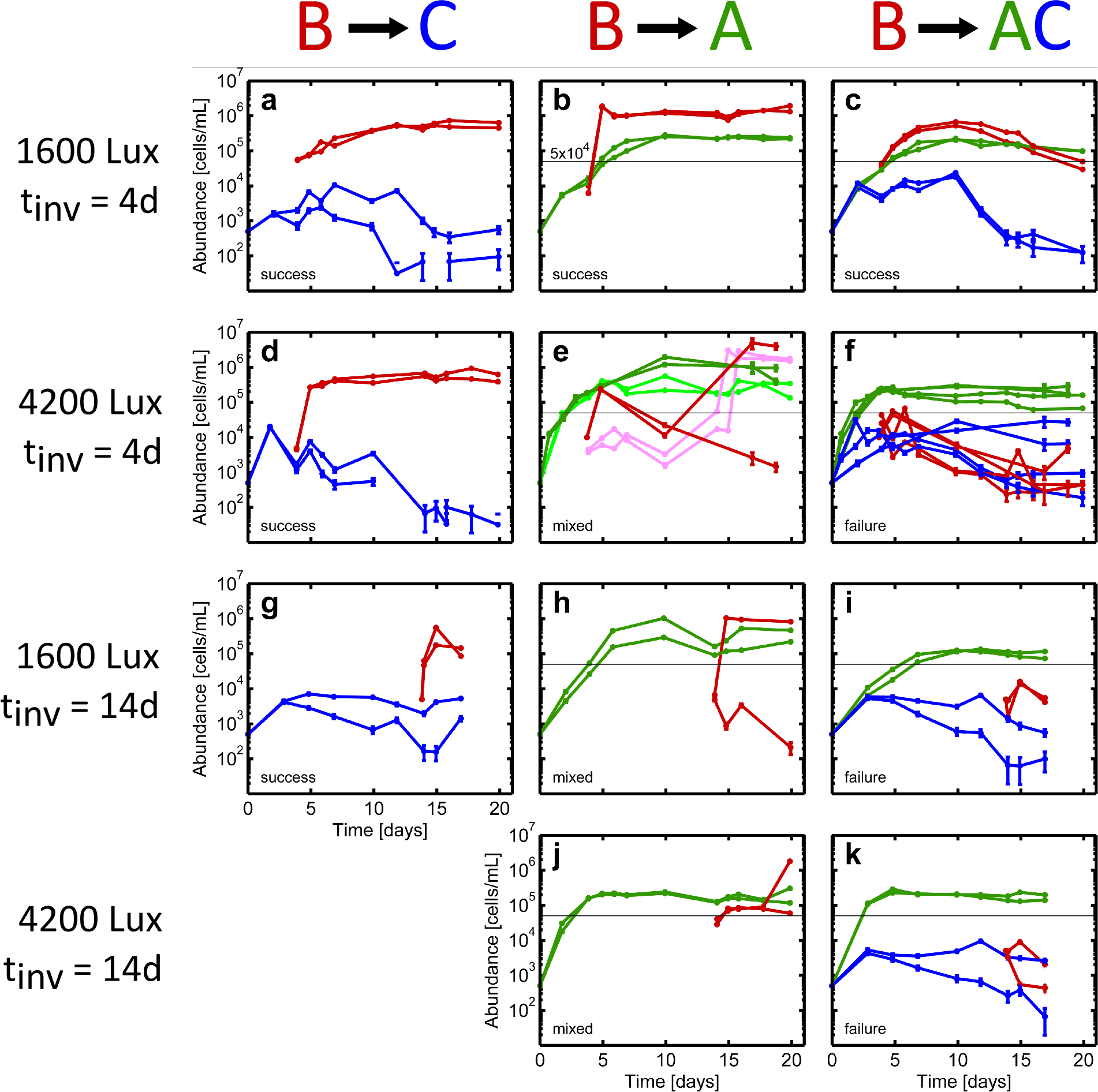
Bacterial invasion dynamics. A, B and C abundances are shown in green, red and blue respectively. B invasions of C (left column) or A (middle column) or AC (right column) occurred in either 1600 Lux (low light) or 4200 Lux (high light) at days 4 or 14 as indicated for each row. In each condition (community composition, light intensity, invasion time) at least two replicate communities were measured. Labels in each panel: “success”, “failure”, or “mixed” indicate the classification of the outcome of bacterial invasion with success defined as B exceeding a density of 7 × 10^4^ mL^−1^ for an extended period of time. Horizontal black line at 5 × 10^4^ mL^−1^ indicates a threshold on A. When A exceeds this threshold, B invasions can be inhibited. In panel (**e**), pink traces for B and light green traces for A denote experiments in which spent media measurements were performed (Fig. S14b). Abundances are measured via flow cytometry. Error bars are computed as described in Methods. For time points where error bars are not visible, errors are smaller than the size of the points. B abundances are reported as the total number of cells including planktonic cells and cells in aggregates as described in the SI.

### Bacterial invasions of algae

Next we studied the dynamics of B invading A (Fig. 2b,e,h,j). When B is introduced into a low-density (<5 × 10^4^ mL^−1^) culture of A, it successfully grows to high density (Fig. 2b). Unexpectedly, when B is introduced into a high density (>5 × 10^4^ mL^−1^) culture of A, B is strongly inhibited, exhibiting either very slow growth eventually reaching high densities more than 7 days after introduction, or crashing to low densities over a period of more than 5 days (Fig. 2e). In these experiments the ultimate outcome of B invasions is variable, with some succeeding and others failing, even between two identically prepared replicates (Fig. 2h,j). We find that inhibition of B growth by A only occurs when A is at high density and that the inhibition occurs both at low and high light levels (Fig. 2e,h,j). When A abundances are high at the time of B introduction, B eventually reaches high density (>7 × 10^4^ mL^−1^) 50 % of the time (5 of 10 invasion experiments, Figs. 2e, 2h, 2j & S9c). However, when A densities are low at the time of B introduction, B grows to high density immediately in every replicate (8 of 8 experiments, Figs. 1f, 2b, S8b & S9b). Note that the inhibitory interaction between A and B is only apparent when B has not reached stationary phase. If B reaches stationary phase before A abundances exceed 5 × 10^4^ mL^−1^, we observe no inhibition of B by A, not even once A abundances exceed that density later in the experiment (compare Fig. 1c&f).

To confirm the dependence of B invasion success on A density, we performed a set of invasion experiments where B was introduced into A cultures at *t* = 0 days (pair-culture experiment), 1 day, 3 days, and 4 days, all at high light (4200 Lux) (Fig. S9). We confirmed that for those A cultures which did exceed density 5 × 10^4^ mL^−1^ at the time of B introduction, B growth was inhibited.

We next showed that resource competition is not the mechanism by which A inhibits B growth. We harvested spent media from an AB invasion experiment at several time points (Fig. 2e pink traces, Fig. S14b). We then filtered out both A and B and inoculated fresh B cells at low density into this spent media. Remarkably, B was able to grow to high density (>10^5^ mL^−1^) on spent media harvested before ~ *t* =15 d. (Fig. S14d). For spent media harvested after ~ *t* =15 d (when B finally grows to high density), B can no longer grow to high density on the spent media. This result shows that consumable nutrients exist for B, even while B’s growth is being inhibited by A. High-density populations of A do not compete with B for these nutrients, but do limit the ability of B to consume them.

### Possible mechanisms of algae inhibition of bacterial growth

We undertook a series of experiments to better understand the mechanism by which A inhibits B growth. We found that the inhibition of B growth by A requires illumination, and that the presence of A alone, in the absence of light, is not sufficient to inhibit B growth (Fig. S10). Literature on algae-bacteria interactions suggests that reactive oxygen species may be responsible for bacterial growth inhibition ^33, 34^, but we showed conclusively that the A was not producing sufficient H_2_O_2_ to limit B growth (Fig. S12). We found that the role of light intensity in the inhibition of B growth was dependent on the growth history of A (Fig. S11). Further, we considered the hypothesis that increased cell-to-cell contact due to the higher density of A could be responsible for the inhibition of B by A. We estimated the frequency of B collisions with A to be extremely high even at low A density and so we expect that an increase in physical contact between A and B due to higher A density would not be responsible for the inhibition (SI). Additionally, detailed analysis of our flow cytometry data showed that A and B stick to each other, but the rate is low (~5 %) and does not change significantly between inhibited and successful invasions (Fig. S13). As a final check, we confirmed that successful invasions were not a case of A detritus being misclassified as B (Fig S19).

Irrespective of the molecular mechanism of the antagonistic interaction between A and B, it is clear that the inhibition of B requires high density A, light, and a B population that has not reached stationary phase. Finally, the variable outcomes for B invading A are in stark contrast to the reproducible dynamics observed in Fig. 1 and in previous studies^25^, suggesting that there may be stochastic processes at the single-cell level which are responsible for the inhibition or that the system is highly susceptible to small experimental variations near the transition between B inhibition and B growth. We now turn to the central objective of the present study, the invasion of B into AC communities.

### Bacteria fail to invade algae-ciliate communities when algal densities are high

Next we performed B invasions of AC cultures (Fig. 2c,f,i,k). Unexpectedly, we found that in every single case where C is present and A density exceeds 5 × 10^4^ mL^−1^ at time of B introduction, B fails to invade and ultimately declines to very low densities (Fig. 2f,i,k). This result stands in stark contrast to B’s successful invasions of A or C alone. When B is introduced into an AC community with low A density (<5 × 10^4^ mL^−1^), B grows to high density and then slowly declines in abundance later in the experiment (Fig. 2c). We conclude that if C is present and A is at high density at the time of B introduction, B cannot proliferate and ultimately declines to low abundance. B’s failure to invade AC cultures is the main finding of this study.

One possible explanation for this finding is that AC communities with high A densities have exhausted a critical nutrient for B growth. Spent media experiments again show this not to be the case. We performed a series of spent media experiments where communities with A, C, or A and C were grown, samples were harvested, and then all cells were removed by filtration (example shown in Fig. S14a). B was then inoculated into this spent media and grown to saturation in a 96-well plate where its abundance was assayed by flow cytometry. This measurement captures the carrying capacity of bacteria on the spent media. We find that B is able to grow on the spent media of A, C, and AC communities to a saturating density that is indistinguishable from growth on fresh media, irrespective of the time at which the spent media was taken (Fig. S14c). This result rules out the hypothesis that nutrient competition accounts for any of the invasion outcomes in Fig. 2.

The results of Fig. 2 suggest that a higher-order effect, unexpected from pairwise interactions, governs the outcome of B invasions of AC communities. Only when A and C are both present do B invasions reliably fail. Next we sought to understand the mechanism of this effect.

### Algae enhance ciliate predation of bacteria in a density-dependent fashion by inhibiting bacterial aggregation

We propose that a higher order (3-body) interaction is responsible for the fact that B cannot invade an AC community when A abundances exceed 5 × 10^4^ mL^−1^: high density A induce B to remain in a planktonic (single-celled) state, robbing B of aggregation, its primary defense mechanism against predation by ciliates (Fig. 1e).

First, we show that B aggregation is inhibited by high density A. Note that as bacteria grow in monoculture they initially aggregate and then ultimately disaggregate (Fig. 1e, orange traces). This non-monotonic change in aggregation is potentially due to substrate level dependent aggregation rates observed previously^35, 36^. When A and B are grown in pair-culture, the aggregation and subsequent dispersal of B is nearly identical to what we observe in B monocultures (compare red traces in Fig. 3e to orange traces in Fig. 1e). Similar B aggregation dynamics are observed when B invades a low density A culture (Fig. 3b,e). However, when B invades at day 4 into a high light (4200 Lux) A culture that has reached 1 × 10^5^ mL^−1^, B does not grow immediately (Fig. 3c) and for the duration of this growth inhibition their aggregation is inhibited (Fig. 3e, black traces). Only once the inhibitory effects of the algae are overcome by B after ten days can B grow and aggregate. We conclude that algae inhibit bacterial aggregation in a manner that depends on the density of algae.

**Figure 3:**
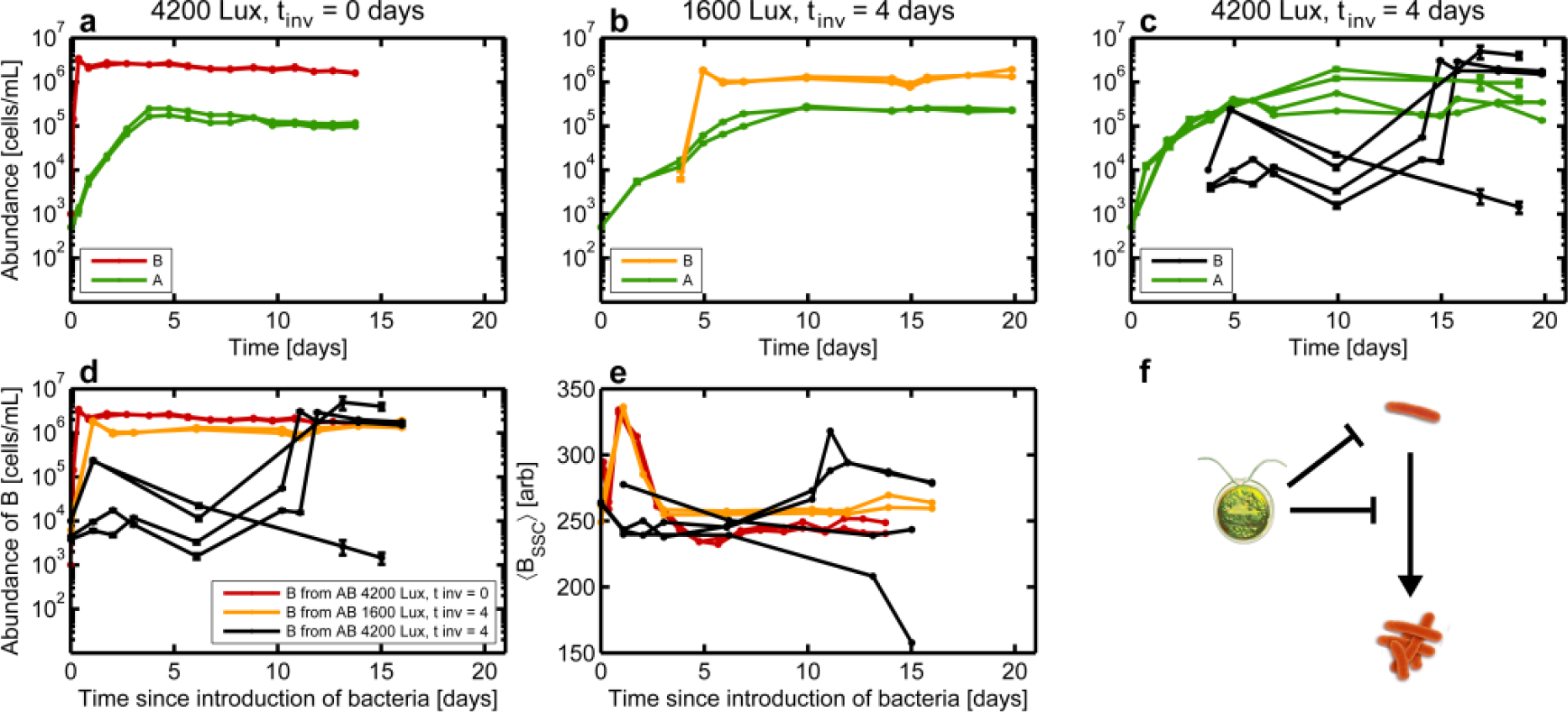
Algae suppresses bacterial aggregation. A abundances are shown in green. Color of B abundance trace varies in **a-c**as indicated in the legends. **a,** Abundance dynamics for two replicates of a high light (4200 Lux) AB pair-culture (t_*inv*_ =0 days) **b,** Abundance dynamics for two replicates of a B invasion on A in low light (1600 Lux) with t_*inv*_ =4 days. A density is ~1 × 10^4^ mL^−1^ at time of B introduction **c,** Abundance dynamics for four replicates of a B invasion on A in high light (4200 Lux) with t_*inv*_ =4 days, A density is > 1 × 10^5^ mL^−1^ at time of B introduction. Abundances are measured via flow cytometry. Error bars are computed as described in Methods. For time points where error bars are not visible, errors are smaller than the size of the points. B abundances are reported as the total number of cells including planktonic cells and cells in aggregates as described in the SI. **d,** Overlay of B abundances from (**a,b&c**), translated so that *t* = 0 corresponds to the time of B introduction. **e,** Overlay of mean side-scatter signal of B, translated so that *t* = 0 corresponds to the time of B introduction. The low side-scatter signal at the final time-point in one of the black traces arises from a small number (20) of counts. Colors of traces in (**d,e**) correspond to panels (**a-c**). **f,** Diagram of interactions for A and B. Panel (**a**) is reproduced from Fig. 1**f**, (**b**) and (**c**) are reproduced from Fig. 2**b&e**.

We investigated the inhibition of B aggregation by A further by examining the effect of A spent media on the ability of B to aggregate (Fig. S15). Using data from the previously described spent medium experiment, we examined the aggregation state of B (mean side-scatter) as a function of the time at which the spent media was extracted from the growing A culture. We find an inverse relationship between the extraction time of spent media and level of B aggregation when grown on that spent media. The fact that A is not physically present in the spent media suggests that its ability to inhibit B aggregation is mediated by a chemical rather than a physical interaction. Interestingly, while spent media from an A monoculture disaggregates B, it does not inhibit growth of B. This result indicates that aggregation is not necessary for B to grow in this media. The independence of aggregation and growth is supported by the fact that a B mutant deficient in aggregation (∆*csgA*) grows in monoculture on this media without aggregating (Fig. S7e). Our observation that A disaggregates B is qualitatively consistent with observations that *Chlamydomonas* can secrete signaling molecules such as auto-inducers which can interfere with biofilm formation^37^.

C has the opposite effect on B in that it induces B to aggregate (Fig. 1e, Figs. S6 and S7). In all cases where only B and C are present, aggregation of B greatly exceeds that in B monoculture or AB pair-culture. We conclude that A and C have opposing effects on the aggregation state of B, with A inhibiting and C enhancing aggregation.

We hypothesize that the failure of B to invade a culture of C and high-density A (Fig. 4c) is due to A inhibiting B aggregation and subsequently increasing the predation pressure of C on B. Under this hypothesis we expect that when B fails to invade an AC community, B will have failed to aggregate, and this is precisely what we observe (Fig. 4e). Conversely, when B successfully invades an AC community (which occurs when A is at low density at time of B introduction) B aggregates effectively as we expect, thus evading predation from C (Fig. 4b,e). We conclude that when A inhibits B aggregation, this results in stronger predation of B by C, thus driving bacterial abundances down in time.

**Figure 4:**
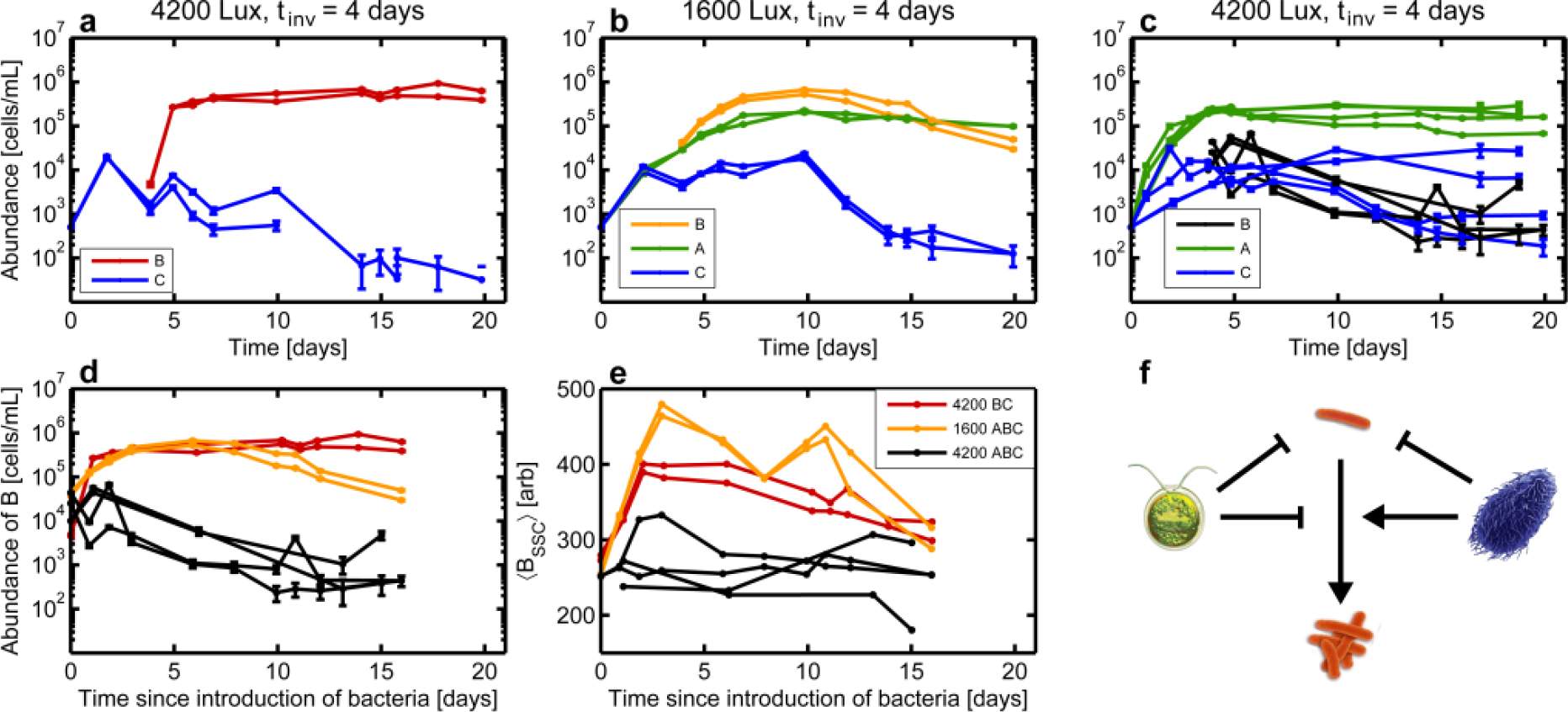
Algae inhibit bacterial aggregation enhancing ciliate predation resulting in invasion failure. A and C abundances are shown in green and blue respectively in all panels. Color of B traces differs between panels **a-c** as shown in the legends. **a,** Abundance dynamics for two replicates of a 4200 Lux (high light) t_*inv*_ =4 days B invasion on a C culture. **b,** Abundance dynamics for two replicates of a 1600 Lux (low light) t_*inv*_ =4 days B invasion on an AC culture. A density is 2 × 10^4^ mL^−1^ at time of B introduction **c,** Abundance dynamics for four replicates of a 4200 Lux (high light) t_*inv*_ =4 days B invasion on an AC culture. A density is > 1 × 10^5^ mL^−1^ at time of B introduction. Abundances are measured via flow cytometry. Error bars are computed as described in Methods. For time points where error bars are not visible, errors are smaller than the size of the points. B abundances are reported as the total number of cells including planktonic cells and cells in aggregates as described in the SI. **d,** Overlay of B abundances from (**a,b,c**), translated so that *t* = 0 corresponds to the time of B introduction. **e,** Overlay of mean side-scatter signal of B, translated so that *t* = 0 corresponds to the time of B introduction. Colors of traces in (**d,e**) correspond to panels (**a-c**). **f,** Diagram of interactions between A, B, and C. Panels (**a,b&c**) are reproduced from Fig. 2d,c&f.

In order for our hypothesis to be true, B aggregation must be a necessary condition for survival of B in the presence of C. To validate this aspect of the hypothesis, we constructed a Δ*csgA* strain of *E. coli* that exhibited dramatically reduced aggregation in liquid culture (Supplementary Figure 11, Laganenka *et al.* ^38^). We introduced this B into a culture of C and hypothesized that it would be unable to invade. Surprisingly, Δ*csgA* aggregated and invaded successfully (Fig. S7). Because it aggregated, this strain did not ultimately test our hypothesis, but it did reveal that there are mechanisms other than curli responsible for aggregation.

Taken together, the results of Figs. 2, 3, & 4 show that a higher-order interaction between the three species governs the outcome of B invasions on AC. In particular, A reduces B aggregation which renders the bacteria susceptible to predation by C. In order to confirm that such an interaction would explain our data and also to generalize our result, we next sought a quantitative model of the dynamics in this community.

### Mathematical model of algae-bacteria-ciliate invasion dynamics

Our objective was to construct a model with a minimal number of free parameters that captures the dynamics we observe in Fig. 2. We chose to construct a purely deterministic model which describes the abundance dynamics of the algae and ciliates (*x_A_*, *x_C_*), non-aggregated bacteria (*x_B_*), aggregated bacteria (*A_B_*), and a single substrate consumed by *x_B_* which we denote *S*. Our model takes the form:

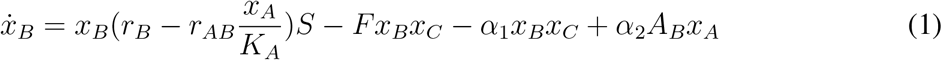

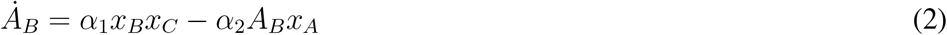

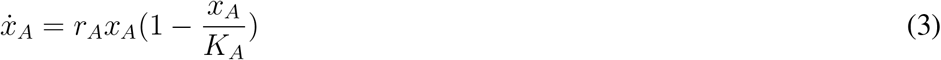

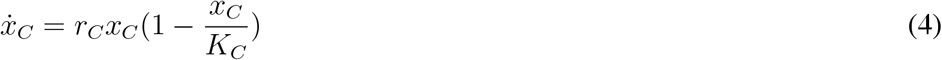

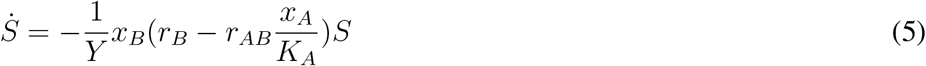

dot denotes a derivative with respect to time, *r_i_* is the growth rate of species *i*, *r_AB_* is a parameter defining the inhibition of *x_B_* by *x_A_*, *F* is a feeding rate, *α*_1_ and *α*_2_ are aggregation and disaggregation rates of bacteria. Note that *A_B_* do not grow (consume *S*) in this model. *K_i_* captures the carrying capacity of species *i* and *Y* is a yield of bacteria growing on *S*.

For a complete discussion of the modeling decisions we made see the SI. Briefly, the first term in Eq. 1 captures the fact that *x_A_* only impacts the growth rate of *x_B_* and not its final abundance. Substrate is considered explicitly to enforce the fact that *x_B_* cannot recover from predation if it had already reached saturating density (such a recovery which would occur with logistic growth). The predation of *x_B_* by *x_C_* is linear in prey density, an assumption justified by the relatively low densities of bacteria in our experiment. We neglect growth of *x_C_* on *x_B_* due to the low yield of ciliates on bacteria and the low densities of bacteria in our experiment. This assumption is supported by the data which shows no substantial difference in C densities with and without B (Fig. S21). Aggregation terms are consistent with our observations in Fig. 1 and 3. In this model, the mechanism by which bacteria (*x_B_* +*A_B_*) fail to invade communities of *x_A_* and *x_C_* is the disaggregation of *A_B_* to *x_B_* in a manner that is dependent on *x_A_* density (*α*_2_*A*_*B*_*x*_*A*_) and the subsequent predation of *x_B_* by *x_C_* (*Fx*_*B*_*x*_*C*_).

The dynamics of *x_A_* and *x_C_* are modeled as logistic growth. For *x_A_*, this model recapitulates the dynamics we observe very well so long as we make *x_A_* growth rate decline as *x_B_* or *x_C_* are added to the community in low light as observed in experiment (Fig. S1a). The model neglects the decline in ciliate abundances at long times, likely due to cell death. All parameters in the model except *r_AB_*, *α*_1_ and *α*_2_ are measured or have been previously reported in the literature (as is the case for *F*). Of these three parameters *r_AB_* must be on the order of *r_B_* if we are to observe any substantial inhibition of *x_B_* growth by *x_A_*. *α*_1_ can be constrained by a close examination of the BC pair-culture dynamics (Fig. S20) and *α*_2_ is treated as a free parameter which we determined by performing a parameter sweep (Figs. S22). With these parameter values (Table S1) we performed numerical integration of the model in Eqns. 1–5 for the invasion experiments shown in Fig. 2. The results are shown in Fig. 5 where we plot total B abundances (*x_B_* +*A_B_*), A abundances (*x_A_*), and C abundances (*x_C_*).

**Figure 5:**
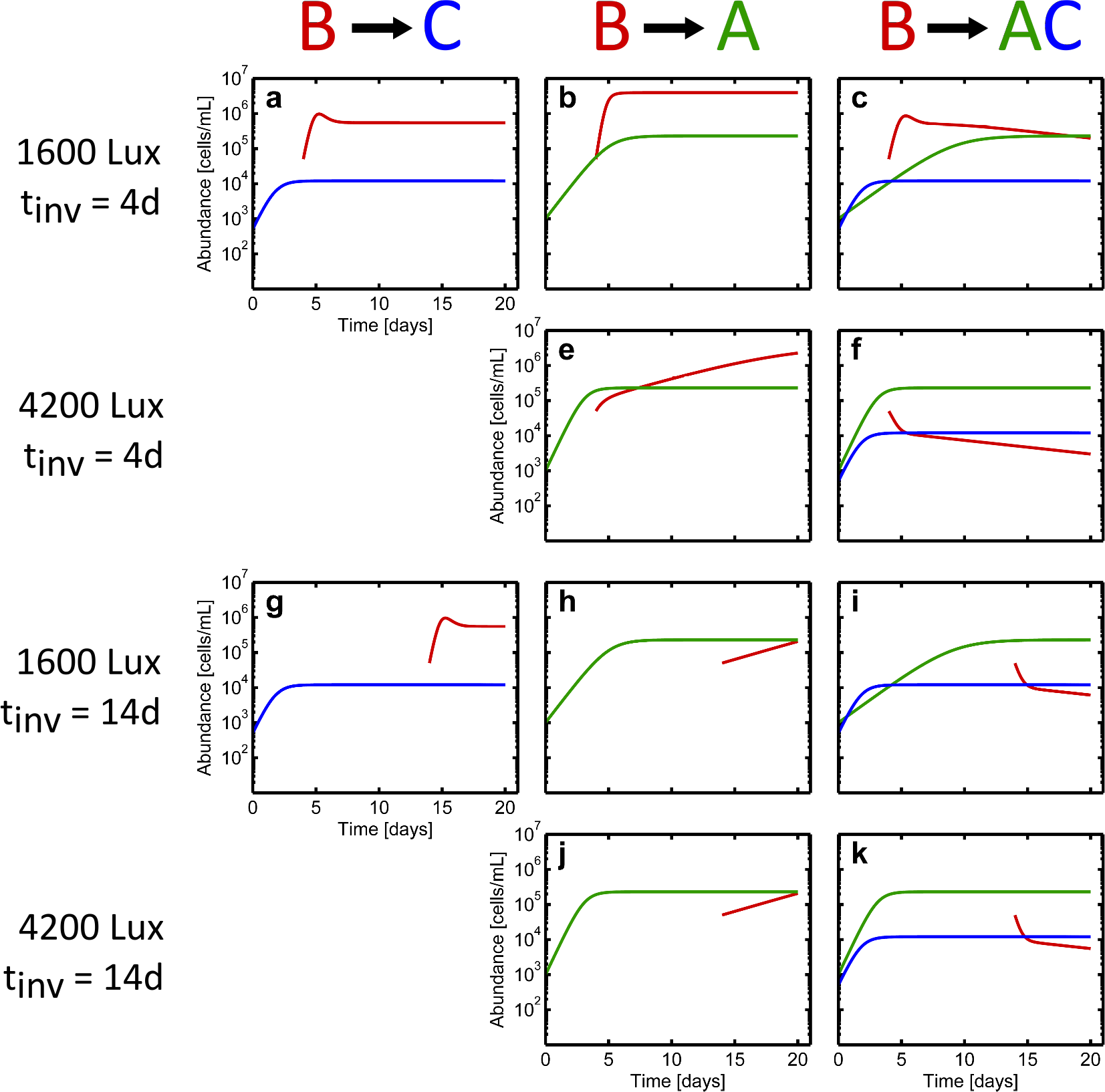
Model of algae-bacteria-ciliate dynamics captures invasion experiment outcomes. Simulation of the model described in the main text. Panels correspond to Fig. 2. Species compositions are organized by column as shown (top). Light conditions and time of invasion are organized by rows as shown (left). In all panels abundance dynamics of A (*x_A_*), B (*x_B_* +*A_B_*), and C (*x_C_*) are shown in green, red and blue respectively. The first panels in the second row and the fourth row are omitted since BC dynamics do not depend on illumination intensity in the model. Parameters of the model are as follows: *r_B_* = 0.3 h^−1^; *r_AB_* = 0.29 h^−1^; *r_C_* = 0.073 h^−1^; *r_A_* (high light) = 0.073 h^−1^; *r_A_* (low light, w/BC) = 0.016 h^−1^; *r_A_* (low light, w/C) = 0.025 h^−1^; *r_A_* (low light, w/B) = 0.031 h^−1^; *r_A_* (low light, alone) = 0.045 h^−1^; *K_A_* = 2.3 × 10^5^ mL^−1^; *K_C_* = 1.2 × 10^4^ mL^−1^; *F* = 1 × 10^−5^ mL h^−1^; *α*_1_ = 2.5 × 10^−6^ mL h^−1^; *α*_2_ = 2 × 10^−8^ mL h^−1^. Substrate concentration is chosen to yield the observed carrying capacity of B in monoculture (3.9 × 10^6^ mL^−1^).

The results in Fig. 5 should be compared to the experiments in Fig. 2. We note that this simple model captures the basic features of the invasion experiments: (i) B successfully invades C cultures (ii) B densities in the presence of C are lower than for B invasions of low-density A cultures (compare Fig. 5a,b) (iii) when B invades a high density A culture, its growth rate is attenuated (Fig. 5e,h,j), but B eventually reaches high density (>1 × 10^6^ mL^−1^) (iv) when B is introduced into an AC culture with high density A, B declines in abundance continuously (Fig. 5f,i,k) (v) when B is introduced into an AC culture with low density A, B invades immediately, but slowly declines in abundance over time (Fig. 5c). Our deterministic model cannot capture the variability in outcomes of B invading A alone. The model describes the invasion experiments faithfully without requiring the specification of a large number of unknown parameters.

### Impact of higher-order interaction is apparent in ABC tri-culture abundance dynamics

We next asked whether the model could provide a non-trivial prediction regarding community dynamics. To investigate, we simulated the dynamics of the ABC tri-culture under the two light regimes. The model predicts that under high light conditions, where *x_A_* rapidly approaches *K_A_* bacterial abundances (*x_B_* +*A_B_*) are attenuated at long times. In contrast, for low light conditions where *x_A_* does not reach high density until the very end of the experiment, our model predicts limited attenuation of bacterial abundances (compare Fig. 6c,d). The lower bacterial abundances observed in our simulation at high light arise from reduced bacterial aggregation and increased predation of *x_B_* by *x_C_* (Fig. 6f).

**Figure 6:**
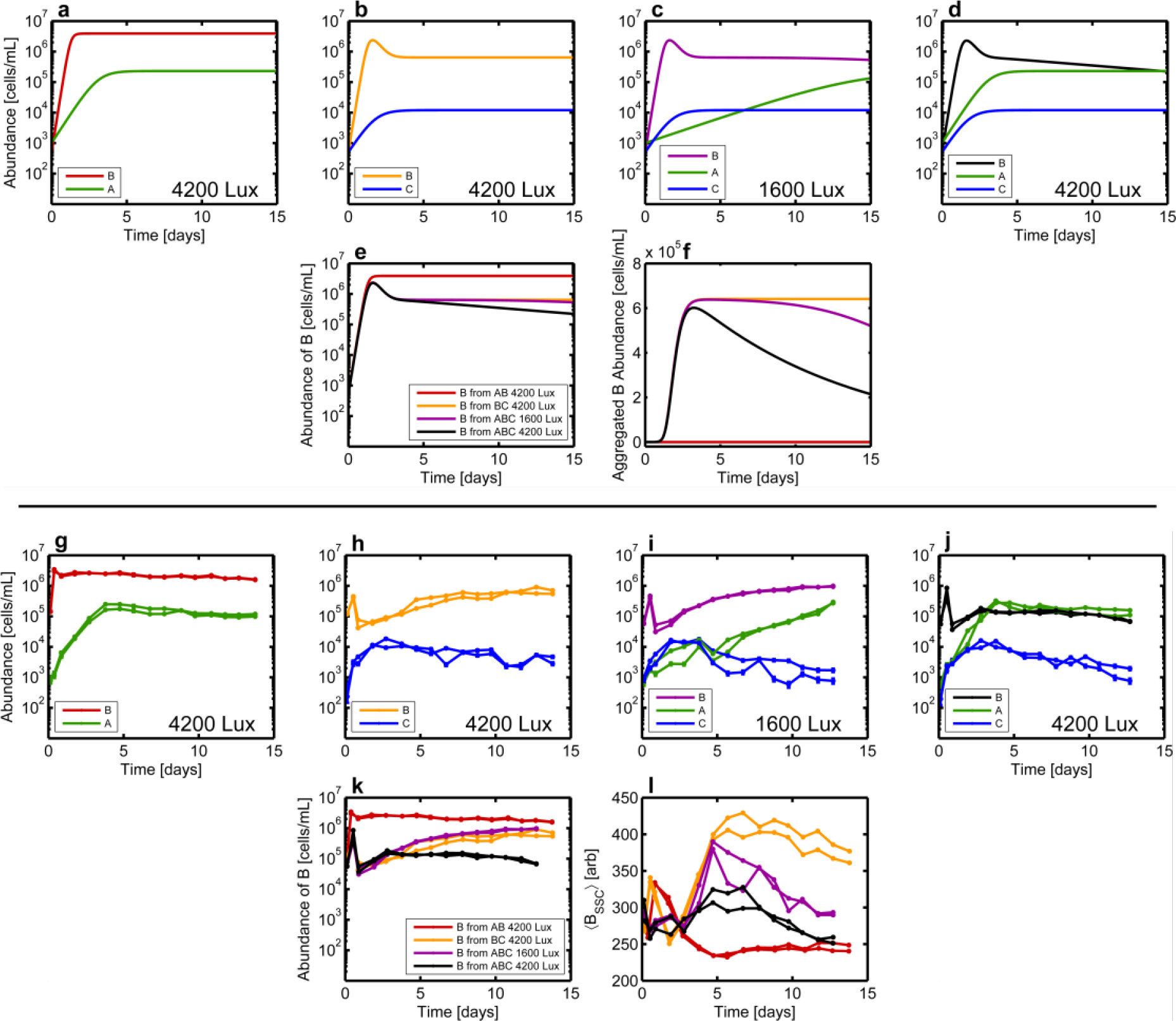
Higher-order interaction impacts algae-bacteria-ciliate tri-culture abundance dynamics. **a-f,** Simulations of abundance dynamics in tri-culture conditions where all species are introduced at low density at *t* = 0 days. All model parameters are given in Table S1. In all panels *x_A_* and *x_C_* are shown in green and blue respectively. The color of B (*x_B_* + *A_B_*) varies as indicated in the legends of (**a-d**) to facilitate the overlay plots in (**e,f)**which show total B abundances (*x_B_* + *A_B_*, **e**) and aggregating cell abundances (*A_B_* only, **f**). Legend from (**e**) applies to (**f**). **g-l**, Experimental measurements of tri-culture abundance dynamics corresponding to panels (**a-f**). Replicate communities are shown in each condition. Abundances are measured via flow cytometry. Error bars are computed as described in Methods. For time points where error bars are not visible, errors are smaller than the size of the points. B abundances are reported as the total number of cells including planktonic cells and cells in aggregates as described in the SI. Note that the color of the traces for B abundances in (**g-j**) correspond to those in (**a-d**) and are indicated in the legends. **k**, Overlay of abundance dynamics for B from (**g-j**). **j**, Mean side-scatter (which is a proxy for level of aggregation) of the B population from panels (**g-j**). Legend from (**k**) applies to (**l**).

To test the predictions of the model we performed tri-culture experiments with the full ABC community at low and high light levels. The dynamics are shown in Fig. 6i,j. In the full ABC ecosystem, the growth rate of A differs dramatically between the two light levels (0.016 h^−1^ for low light versus 0.073 h^−1^ for high light). The slow growth rate of A in low light results in A reaching saturation only after 14 d. In contrast, in high light, A reaches saturating densities in 4 days. When we compare the dynamics of B in the full ABC ecosystem in low and high light we observe that high light results in a substantial decline in bacterial abundances by the end of the experiment (Fig. 6k, purple and black traces). In contrast, when B is grown with only C (and not A) in high light there is no decline in B abundances at long times (Fig. 6k, orange and purple traces). When we examine the bacterial aggregation state in the ABC low light and high light conditions we find less B aggregation in the high light condition (Fig. 6l). We therefore conclude that in the high light condition the rapid rise of A drives substantial disaggregation of B and subsequent predation of B by C. Note that the effect is not driven by light alone since changing light levels does not alter the abundance dynamics of B in monoculture, AB pair-culture or BC pair-culture (Fig. S8). The predictions of the model are confirmed and so we conclude that rapid growth of A to high density substantially impacts B in the presence of C via the same higher order interaction which inhibits B invasion of AC communities with high A abundances.

## Discussion

Our results show that phototrophs can indirectly inhibit the growth of heterotrophic bacteria through higher order interactions with predators. Phototroph-heterotroph interactions are known to be mediated by competition and cross feeding^39^, but our data provide a new mechanism by which phototrophs might keep faster growing heterotrophs from consuming available nutrients such as nitrogen or phosphorous.

In the context of the invasion literature (recently summarized be Mallon *et al.*^20^) our results show that resource competition alone is insufficient for predicting the outcome of invasions in communities where antagonistic interactions are present. The result has important implications for understanding community structure from coral reefs to wastewater treatment facilities where such interactions are known to be present^18, 19^.

Further, recent theoretical work suggests that higher-order interactions (3-way and higher) enhance the stability of complex communities whereas communities described by pairwise interactions alone become less stable as complexity increases^40, 41^. While several studies have previously detected the presence of higher-order interactions^42, 43^, direct measurements of the impact of higher-order interactions on ecosystem dynamics or invasions is limited. In contrast, several studies of dynamics in communities where interactions are mediated by resource competition have shown that pairwise interactions are sufficient to describe community dynamics^44, 45^. Our study shows definitively that for a model community which includes predation and little or no competition for resources, higher order interactions are not only present, but have substantial impacts on dynamics (Fig. 6). To move theory closer to observation and experiment, it will be critical to investigate whether higher-order interactions are more common when antagonism is present or not.

Two previous studies on the ABC community, or a closely related ecosystem with *C. reinhardtii* replaced with *Euglena gracilis*, under hermetically sealed conditions have looked at interactions between these species. One study compared interactions inferred from abundance fluctuations in the full three species community to interactions measured via pair-culture experiments^46^. A second study also used pairwise experiments^47^. In the case of the pairwise experiments, both studies detected evidence for the antagonistic effect of A and C on B. Interestingly, inference of pairwise interactions from abundance fluctuations in an ensemble of ABC communities did not identify these negative interactions^46^, and instead inferred positive effects between all three species. This result points to the challenge of inferring interactions from fluctuations in time series data^48–50^. Inferring interactions from fluctuations should work well near a fixed point of the community dynamics. However, understanding the full non-linear dynamics governing the community is likely a necessity for predicting the outcome of processes like invasion where abundances can change by several orders of magnitude. Finally, neither previous study of the ABC system made a statement about the presence of higher-order effects in the community or the mechanisms mediating these interactions.

More broadly, our study provides a stark example of the level of complexity present in even a comparatively simple model microbial community. Our discovery that a phenotypic trait of the bacteria (aggregation) is modulated by the algae and dramatically impacts the success of bacterial invasion points to both the power of studying model communities like the ABC system and the challenge of predicting dynamics in more complex ecosystems.

## Methods

### Strains

The algae is *C. reinhardtii*, strain UTEX2244 obtained from the UT Austin Culture Collection of Algae `utex.org`. All *C. reinhardtii* cells are of a single mating type mt+ and grow vegetatively. Algae are cryogenically preserved and stored in a liquid nitrogen https://utex.org/pages/cryopreservation#liquid. The *E.coli* strain is MG1655 Δ *flu*, Δ *fimA* and was constructed previously ^24^. A constitutively expressed yellow fluorescent protein (YFP, promoter *λ*P_*R*_) was transduced into the genome with phage P1. The donor strain for YFP fluorescence has the YFP gene inserted in the *intC* locus along with a chloramphenicol antibiotic resistance marker ^51^. The Δ*csgA* strain was constructed by P1 transduction from KEIO collection^52^ mutant into MG1655 (WT) background. The same YFP construct was also transduced into this strain. The ciliate is *T. thermophila*, strain CU428.2 obtained from the Cornell University Tetrahymena Stock Center https://tetrahymena.vet.cornell.edu/. All *T. thermophila* cells are of mating type VII so there is only asexual reproduction. This strain grows vegetatively indefinitely without sexual reproduction. Ciliates were cryogenically frozen and stored in a liquid nitrogen ^53^.

### Culturing

Before beginning the experiment, each of the organisms is cultured separately in distinct media. *A* is cultured in a 30 °C shaker-incubator with ~3000 Lux illumination in TrisAcetate-Phosphate (TAP) media inoculated directly from a freezer stock. Algae used in co-culture experiments (low-light), A, AC, ABC and (high light) A, AC and ABC were grown at 25 °C prior to the experiment. TAP is a defined media with acetic acid as a carbon source https://www.chlamycollection.org/methods/media-recipes/tap-and-tris-minimal/. *C* is cultured in a 30 °C stationary incubator in (undefined) SPP media inoculated directly from a freezer stock. *B* was cultured in a 30 °C shaker-incubator in 1/2x Taub 0.03% proteose peptone No. 3 inoculated from a single colony grown on an lysogeny broth (LB) plate.

### Control of initial conditions

The cultures of each of the three organisms are washed twice into 1/2x Taub .01% Proteose Peptone No. 3. Flow cytometry is performed on a sample from each washed culture to estimate cell densities. The cultures are then diluted into 1/2x Taub .01% proteose peptone No. 3 in order to achieve nominal densities of 500 ± 22 mL^−1^ for *A*, 1000 ± 32 mL^−1^ for *B*, and 500 ± 22 mL^−1^ for *C*. Error bars are assumed from Poisson counting error. Organisms are always started at these densities at the beginning of an experiment, regardless of whether that experiment is monoculture, pair-culture, or tri-culture. For B invasion experiments the starting density was 1 × 10^4^ mL^−1^.

### Experimental conditions

All experiments are performed in 1/2x Taub .01% proteose peptone No. 3. This media is used because it is similar to media used in previous studies with the ABC community^24, 25^ and because each of the three organisms can grow on this media in monoculture, pair-culture, and tri-culture. Taub media is a freshwater mimic media that was originally created to support co-cultures of *Daphnia pulex* and *Chlorella pyrenoidosa* ^54, 55^. It contains 15*µ*M H_3_BO_3_; 0.5*µ*M ZnSO_4_; 3.5*µ*M MnCl_2_; 0.5*µ*M Na_2_MoO_4_; 0.1*µ*M CuSO_4_; 0.5*µ*M Co(NO_3_)_2_; 100*µ*M MgSO_4_; 100*µ*M KH_2_PO_4_; 5.6*µ*M EDTA; 5.6*µ*M FeSO_4_; 1.5mM NaCl; and 1mM CaCl_2_. Proteose peptone No. 3 is an undefined nutrient source that is an enzymatic digest of protein and supplies nitrogen and carbon [http://www.bdbiosciences.com/ds/ab/others/Proteose_Peptone_No_2_3_4.pdf]. The media is titrated to pH 7 before use.

### Culture devices and conditions

During the experiment, 30 mL of culture are grown in a glass vial (Chemglass CG-4902-08 40 mL volume). A 0.1 µm filter allows gas exchange between the culture and the atmosphere. We expect this gas exchange (venting) coupled with stirring allows the community to be rapidly equilibrated with atmospheric O_2_ and CO_2_ concentrations. Eight vials are run in parallel.

Each of the eight vials are kept in an experimental apparatus for the duration of the experiment. The vial fits snugly into a metal block that is held at 30 °C via PID control. Temperature is measured by a thermometer embedded in the metal block and heating/cooling is performed by a Peltier element^35^. The temperature within a vial fluctuates with standard deviation 0.02 °C as determined by the feedback thermometer embedded in the metal block housing the vial. The temperature across the eight vials varies with standard deviation 0.08 °C as measured in a control experiment where each vial is filled with water and the temperature is measured using a high-accuracy Fisher Scientific Traceable Thermometer (p/n: 15-081-102) and taking the standard deviation across vials.

The vial is illuminated by a single LED (Cree XLamp XP-E2 Single 1 Up Neutral White 4000 K color temperature, LED Supply p/n: CREEXPE2-740-1) from below. The LED is driven by an LED driver (BuckPuck DC LED Driver LED Supply p/n: 03021-D-E-350) and the intensity of these LEDs was found to be too high for bacterial growth and is decreased through the use of a neutral density filter. The illuminance is further modulated by applying a voltage to the control pin of the LED Driver. Experiments are performed at either low light (1600 ± 140 Lux) or high light (4200 ± 330 Lux). These values represent the time-averaged illuminance an organism would experience assuming it spends an equal amount of time at each height in the vial. These values are calculated based on measurements of illuminance taken from the top of the metal block with a light meter (LED Light Meter p/n: PCE-LED 20). Error bars are standard deviation across systems. The light levels are on the same order of magnitude as those used in a previous study of the ABC ecosystem^25^. The experimental apparatus is mounted on a stir-plate (Thermo Scientific Cimarec-i Mono Direct Stirrer 50095601) that keeps cultures stirred at 444 ± 4 RPM. This rotation speed was measured with a custom Hall Probe. Error is standard deviation across systems.

### Sampling of communities for flow cytometry

Samples of communities are taken via a syringe attached to a sterile port. 500 µL are drawn from the vial for each sample. This process constitutes destructive sampling of the community. Over the course of a typical experiment, 16 samples are taken from the vial which corresponds to 8 mL being removed from the vial. The depth of liquid in the vial decreases by 1.65 cm from its initial depth of 6.25 cm.

### Abundance measurement by flow cytometry

Flow cytometry is performed using a Becton-Dickson LSR II. To count bacteria, YFP fluorescence is plotted versus side-scatter (SSC) and cells are gated manually. To count algae, Chlorophyll-b fluorescence is plotted versus YFP fluorescence and gated manually. To count ciliates, CFP fluorescence is plotted versus SSC and the gate is drawn manually. Correct gating is confirmed by making measurements on monocultures. Because of the size difference between bacteria and ciliates, the two organisms scatter vastly different amounts of light and so a different gain is appropriate for the SSC channel for each. We therefore run every flow sample on two different settings. Settings 1 is used for ciliates and gain voltages are 498 for CFP and 203 for SSC. Settings 2 is used for algae and bacteria and gain voltages are 501 for YFP, 275 for chlorophyll-b, and 250 for SSC.

### Calibration of flow rate to infer densities

To report densities we calibrate the liquid flow rate through the flow cytometer using Spherotech Accucount fluorescent beads (ACFP-50-5, 5.0-5.9 *µ*m) which come at a known concentration of 2 × 10^6^ mL^−1^. The beads are diluted tenfold and run for 30 s, the same duration of time that ecosystem samples are run. From the number of beads detected in 30 s we compute a flow rate. For every time point in the abundance data, we perform three replicates of this volume calibration. We assume that the volume calibration applies to all samples run at that time point (within ~1 hour of the calibration of the cytometer). Over the course of this study (2 years) we observed a decrease in the LSR II flow rate by 60%.

### Error bars on abundance measurements

Error bars on abundances are calculated by performing propagation of two forms of error: (1) the Poisson error inherent in counting a finite number of cells and (2) the uncertainty in volume run through the flow cytometer. The uncertainty in flow rate is taken as the standard deviation across the three replicates of bead calibration on the day the abundance was measured.

### Spent media experiments

In spent media experiments, cultures are prepared and grown as normal. To obtain spent media, several hundred microliters are extracted from the culture and filtered through a 0.22 µm polyethersulfone (PES) filter. Spent media is stored at 4 °C until all spent media extractions are complete. Spent media was then added to a 96-well microtiter plate and inoculated with bacteria and, depending on the experiment, ciliates. The plate was shaken and incubated at 30 °C for two or three days so that bacteria can grow to saturation. Bacterial abundance is then measured via flow cytometry.

### Simulations

Numerical integration of the model was performed using custom written Matlab scripts. Time steps of 1 minute were used to ensure numerical stability and accuracy. Organisms that fell below a density of 1 mL^−1^ were assumed extinct and could not recover. Invasions were accomplished by instantaneously adding a fixed density of B at *t_inv_*. Parameters of the model and details of the modeling choices are available in the SI.

## Acknowledgements

The authors acknowledge Jason Merritt, Karna Gowda, Barbara Pilas, and Elizabeth Ujhelyi. Additionally, the authors acknowledge support from the Department of Physics, the Carl R. Woese Institute for Genomic Biology, and the Roy J. Carver Biotechnology Center, all at the University of Illinois at Urbana-Champaign. The authors thank Aaron Bell for permission to use the image of the ciliate.

## Correspondence

Correspondence and requests for materials should be addressed to S.K.

