## Supplementary Information for "Higher-order interaction inhibits bacterial invasion of a phototroph-predator microbial community"

### Supplementary material for *Higher-order interaction inhibits bacterial invasion of a phototroph-predator microbial community*

Harry Mickalide & Seppe Kuehn

**Contents**

**Algae and Ciliate growth rates** 1

**Bacterial aggregation** 2
Vortex experiment confirms side-scatter measures aggregation 2
Construction of bacterial aggregate correction algorithm 3
Ciliates cause bacteria to aggregate 7
Ciliate induces aggregation in a $\Delta$*csgA E. coli* mutant 7

**Light level does not affect bacterial or ciliate abundance dynamics** 7

**Algae-bacteria interactions** 8
When algae is at sufficiently high density, it stochastically prevents bacterial invasion 8
Algae must be physically present and illuminated to inhibit bacterial invasion 9
There exists a threshold light level below which algae cannot suppress bacterial invasion 11
Hydrogen peroxide is not responsible for A inhibiting B invasion 11
Algal-bacterial adhesion and invasion supression 12
Physical collisions between bacteria and algae are frequent even when algal density is low 13

**Spent Media Experiments** 14
Neither algae nor ciliates compete with bacteria for nutrients 14
Algae spent media de-aggregates bacteria 15
Ciliates present in algal spent media is not sufficient to prevent bacterial invasion 16

**Live-dead staining experiments** 16

**Successful bacterial invasions are not an artifact of mis-classifing algae detritus as bacteria** 18

**Details of model of ABC dynamics with higher order interactions** 19
Bacteria-ciliate interactions 21
Predation rates and functional form 22
Algae-bacteria interactions 23
Aspects of the dynamics not captured by the model 24

**References** 26

#### ALGAE AND CILIATE GROWTH RATES

We calculated growth rates for algae and ciliates in all coculture experiments. Growth rate was calculated by fitting a line to the linear portion of the natural logarithm of that species' abundance in time. Algae growth rates at 1600 Lux (low light) were significantly different for different species compositions and we thus report a growth rate for each species composition (Fig. S1a). For algae at 4200 Lux (high light), or ciliates at either light level, growth rates were not significantly different across species compositions and we thus report a single growth rate over all species compositions (Fig. S1b,c&d). We do not report growth rates here for bacteria since the time resolution of our flow cytometry measurement is too coarse to reliably measure bacterial growth rates. Bacterial growth rate is instead measured using continuous absorbance measurements in a plate reader.

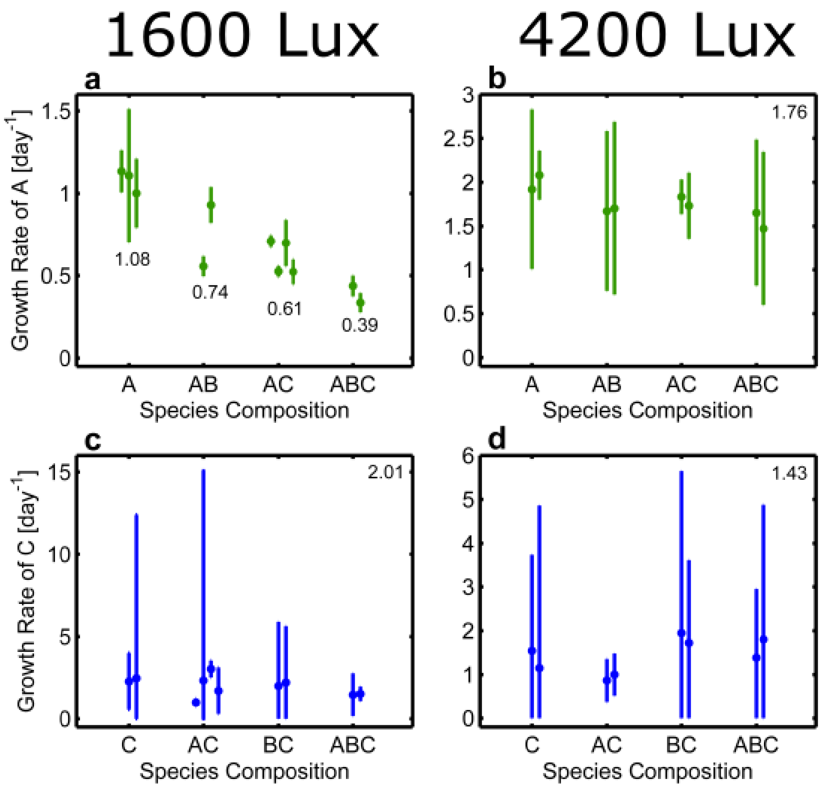


FIG. S1: **Algae and Ciliate growth rates a,** Growth rate of algae plotted for all relevant species compositions for experiments performed at 1600 Lux (low light). The growth rate was calculated for each replicate. Error bars are 95 % confidence intervals. Numbers reported are the mean growth rate across replicates for that species composition. **b,** Growth rate of algae plotted for all relevant species compositions for experiments performed at 4200 Lux (high light). Number reported in the top right represents the mean growth rate across all replicates of all species compositions. **c,** Growth rate of ciliates plotted for all relevant species compositions for experiments performed at 1600 Lux (low light). Number reported in the top right represents the mean growth rate across all replicates of all species compositions. **d,** Growth rate of ciliates plotted for all relevant species compositions for experiments performed at 4200 Lux (high light). Number reported in the top right represents the mean growth rate across all replicates of all species compositions.

#### BACTERIAL AGGREGATION

In the main text we report that side scatter signal of bacteria (YFP fluorescence) reflects the aggregation of bacteria, with higher side scatter levels indicating larger bacterial aggregates. Here we support this claim experimentally.

##### Vortex experiment confirms side-scatter measures aggregation

High side-scatter bacterial objects were originally suspected to be aggregates of bacteria for two reasons. 1) Side-scatter is a measure of how much light is scattered at a 90 degree angle when an object passes through the flow cytometer and larger objects tend to scatter more light. Indeed the ciliates, the largest of the three organisms, has the highest side-scatter signal. Since aggregates of bacteria are larger than single cells, they should have higher side-scatter signal. 2) The high side-scatter portion of the bacterial population stays present throughout an experiment only when ciliates are present and ciliates are known to induce bacteria to aggregate (see main text for discussion).

We performed a vortexing experiment to test if high side-scatter signal bacterial objects were indeed aggregates. In the experiment we performed flow cytometry on a sample of bacteria (Fig. S2a-d), vortexed the sample, and then performed flow cytometry again (Fig. S2e-h). We hypothesized that vortexing would break up aggregates. This hypothesis lead to two clear predictions: (1) that the number of bacterial objects would increase, due to aggregates being broken up into multiple objects, and (2) that vortexing would reduce the number of high side-scatter objects. Both predictions were confirmed. By comparing the bacterial object abundances from before and after vortexing, one can see that abundances increased after vortexing (compare densities reported in upper left corner of Fig. S2a-d to

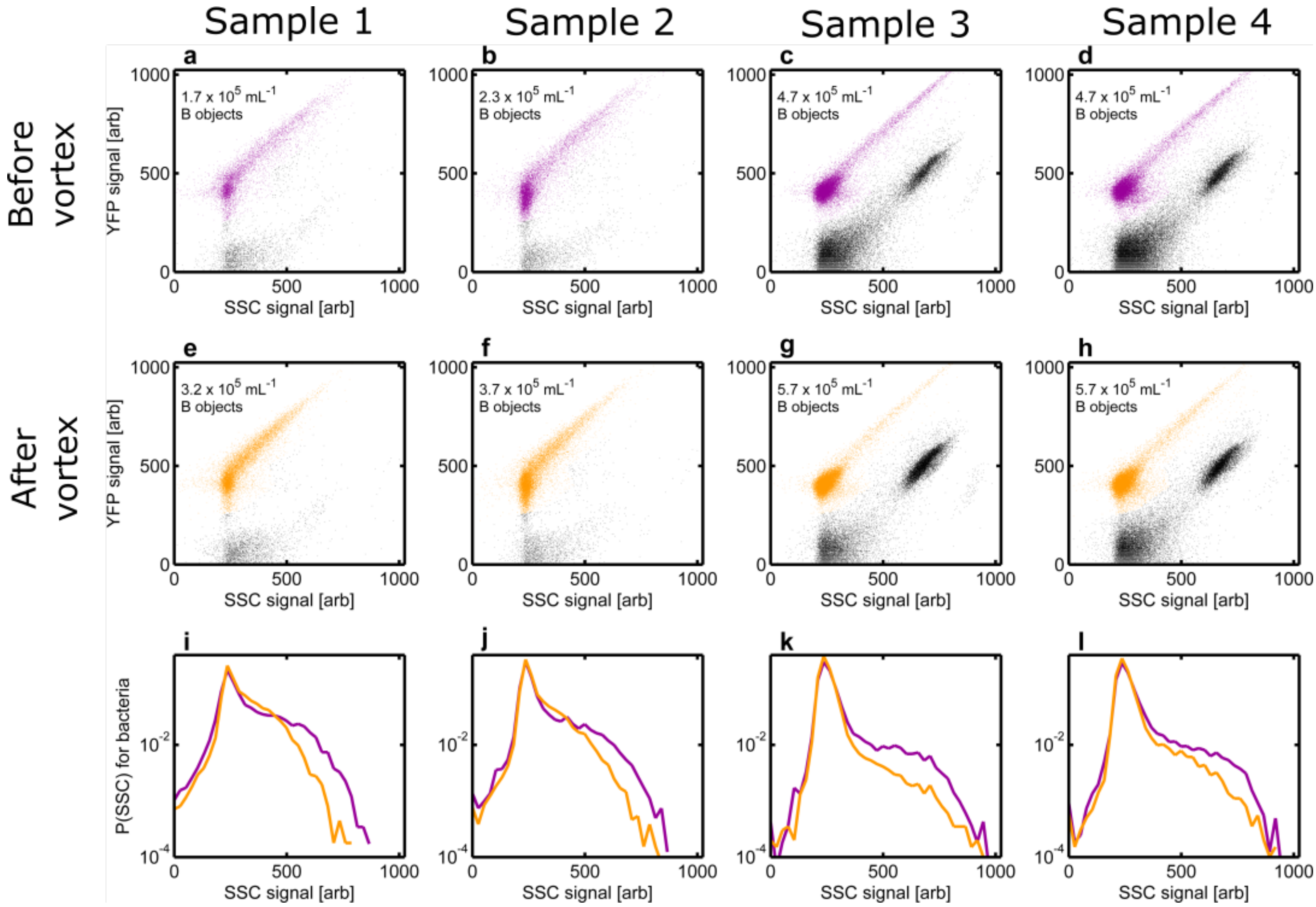


FIG. S2: **Vortex experiments shows high SSC objects are bacterial aggregates a-d,** Yellow fluorescence signal (YFP) plotted versus side-scatter signal (SSC) for flow cytometry data from day 12 of a 1600 Lux BC coculture (**a,b**) and a 1600 Lux ABC coculture (**c,d**). Points colored purple are classified as bacterial objects. The number reported in the plot indicates abundance of bacterial objects. **e-h**, Flow cytometry data from those same samples, but after vortexing. Points colored yellow are classified as bacterial objects. In all cases the abundance of objects classified as bacteria increases after vortexing. **i-j,** Overlays of histograms of side-scatter signal of bacterial objects from before and after vortexing.

panels e-h). By comparing the histograms of side scatter signal of bacterial objects (Fig. S2i-l), one can see that the number of high side-scatter bacterial objects decreased after vortexing. In additional, one can see that vortexing does not affect the location of the peak of the histogram, which therefore presumably corresponds to single-celled bacteria and which are not disrupted by vortexing.

#### Construction of bacterial aggregate correction algorithm

In order to estimate the true number of bacterial cells, we devised a technique to estimate the number of bacterial cells in an aggregate. Having confirmed in the previous section that side-scatter signal correlates with the number of bacteria in an aggregate, we seek an expression for the number of bacteria ($N$) as a function of the magnitude of the side scatter signal for each object ($S$). We assume that for objects below a threshold value of $S < t$ then $N(S) = 1$. We set $t$ to a value which corresponds to the right shoulder of the left mode of the distribution in Fig. S3b. For aggregates, that is objects with $S > t$, we take the ansatz that $N(S) = \alpha S^{\beta}$

$\beta$ is an exponent which represents how the number of cells in an aggregate scales with the side-scatter signal and $\alpha$ is a prefactor. We chose this form because it is monotonic with $S$, it is possible to infer these parameters from our data and scattering theory for simple objects (e.g. spheres) shows that the scaling of scattered light intensity is polynomial in particle size.

The first step in determining $\alpha$ and $\beta$ is to define $t$. $t$ should be the value of side-scatter past which bacterial objects are predominantly aggregates. We estimate this value by plotting a histogram of side-scatter signal of bacteria and marking the point at which the approximately Gaussian curve (representing single cells) turns into a tail (Fig. S3b). Based on visual inspection of histograms of side-scatter, we set $t = 300$.

In all flow cytometry data reported up to this point, what is reported is the *log* of the fluorescence or scattered intensity. When data files of the format used in this study (fcs2.0) are exported from the flow cytometer, all fluorescence/scattering intensities are given as integers on a scale from 0 to 1023 (10-bit ADC), where 0 represents no signal and 1023 represents a signal which saturates the photomultiplier tube. The user is given the option of whether they want the data to be log-transformed or not. Our data are log-transformed since this affords us a larger dynamic range. Therefore, our flow data report $log(S)$ rather than $S$. In order to follow our ansatz above it is necessary to transform our flow cytometry data from a logarithmic to a linear scale. However, the coefficients of the exponential transformation are not given by the manufacturer of the instrument, so it was necessary to infer the parameters of this transformation. To accomplish this we exported a single dataset on a linear and log scale (e.g. $S$ and $log(S)$). From these two datasets we inferred that:

$$S = 0.0899e^{.0092log(S)}$$

(Fig. S3c). Therefore, our threshold on $log(S)$ of 300 is 1.41 on a linear scale.

We next inferred $\alpha$ and $\beta$ from the vortexing experiment shown in Fig. S2. The key insight is that the *total* number of bacterial cells cannot change due to vortexing (although the number of detected objects does change due to vortexing disrupting aggregates). We employed the following method to determine $\alpha$ and $\beta$.

For a given run of the flow cytometer consider the $M$ objects which are detected and classified as bacteria to be indexed by $i$. The side scatter signal from the $i$th object is then $S_i$ which contains a number of bacterial cells $N(S_i)$. For sample $k$ we denote the side scatter signals from all $M$ objects *prior* to vortexing as $S_{i,p,k}$. For the same sample we refer to the side scatter for all objects *after* vortexing as $S_{i,v,k}$ where $i$ now runs to $M'$ with $M < M'$. For example, the distribution of $log(S_{i,p,k})$ is given by the purple traces in the bottom row of panels in Fig. S2 and the distribution of $log(S_{i,v,k})$ by the yellow traces.

Under the assumption that the number of cells (not objects) cannot change due to vortexing the following equality must hold:

$$\sum_i^M N(S_{i,p,k}) = \sum_i^{M'} N(S_{i,v,k})$$

from this we find that:

$$q = \frac{\sum_i^M \alpha(S_{i,p,k})^\beta}{\sum_i^{M'} \alpha(S_{i,v,k})^\beta} = 1$$

Our objective then is to determine the values for $\alpha$ and $\beta$ such that $q = 1$. We now consider $q(\alpha, \beta, k)$ which shows how the ratio of the number of inferred bacterial cells depends on $\alpha$ and $\beta$. A heatmap of $q(\alpha, \beta, 1)$ is shown in Fig. S3d with the important modification that we plot $1/q$ for values of $q < 1$. Local minima in this heatmap near 1 reveal values of $\alpha$ and $\beta$ where our assumptions are satisfied. Note how the pairs of $\alpha$ and $\beta$ along the ascending diagonal of this heatmap have $q \approx 1$. We now compute the same heatmap for $k = [1, 2, 3, 4]$ and compute

$$\sum_{k=1}^4 q(\alpha, \beta, k)^2$$

again taking $1/q$ when $q < 1$ (Fig. S3e). This plot reveals a range of both $\alpha$ and $\beta$ for which this sum is 4 where our assumptions are satisfied for all four samples in our vortex experiment. We applied the square to each element in the sum to especially penalize samples which had high $q$ at the given $\alpha$ and $\beta$.

To narrow the range of parameters we apply a final criterion. Given our assumption that the lower mode of the distribution of $S$ comes from single cells, we know that $N(S < 1.41) \approx 1$. Therefore we computed $N(S = 1.41)$ as a function of $\alpha$ and $\beta$ and the result is shown in Fig. S3f. These criteria alone do not uniquely determine $\alpha$ and $\beta$ so we proceed by selecting $\alpha = 0.9$ and $\beta = 0.7$ where $N(S = 1.41) = 1.14$. This decision is subjective, but does not dramatically alter our results. For all bacterial abundances reported in the main text we use this aggregate correction algorithm.

The success of the aggregation correction algorithm can be seen in how it eliminates a spurious drop in bacterial abundances that was caused by aggregation. In Fig. S3h, we have plotted an abundance curve for bacteria in a

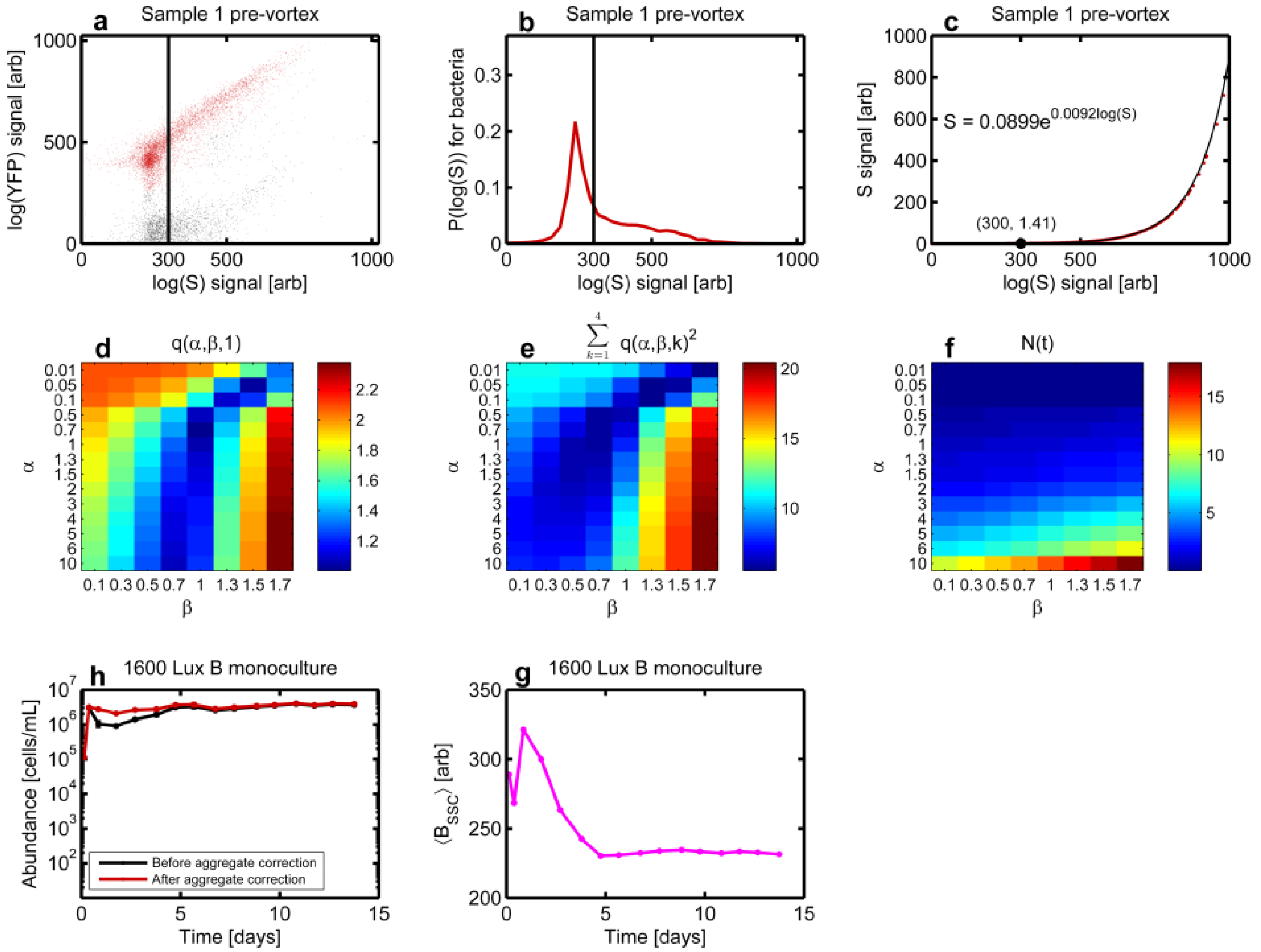


FIG. S3: **Constructing the aggregate correction algorithm a,** YFP plotted versus SSC for flow cytometry data for sample 1 prior to vortexing (Fig. S2a). Red points indicate objects we have labeled bacteria. The black line indicates the threshold between single cells and aggregates at $t = 300$. **b,** Histogram of $log(S_{i,p,1})$ signal for bacteria with the threshold indicated by the line. **c,** Plotting $S_{i,p,1}$ vs $log(S_{i,p,1})$ (red) with the fitted curve (black). The threshold is indicated by the black labeled point. **d,** A heat map of $q(\alpha, \beta, 1)$ where we have plotted $1/q$ when $q < 1$. **e,** A heatmap of $\sum_{k=1}^{4} q(\alpha, \beta, k)^2$ where again we take $1/q$ for values of $\alpha$ and $\beta$ where $q < 1$. **f,** $N(S = 1.41)$ as a function of $\alpha$ and $\beta$ (recall $t = 1.41$). **h,** Abundance dynamics for a 1600 Lux (low light) B monoculture before (black) and after (red) aggregate correction. **g,** Mean SSC (aggregation) plotted versus time for that same 1600 Lux B monoculture.

1600 Lux (low light) monoculture before and after the aggregate correction algorithm is applied. Before correction, all bacterial objects are weighted equally, meaning that an aggregate and a single cell are both counted as a single bacterium. This equal weighting leads to a spurious fall in bacterial abundance after reaching the initial peak. Notice how the apparent decline in bacterial abundance at day 1 corresponds in time to an increased level of aggregation (Fig. S3g). Once the bacteria eventually disaggregate, around day 5 or so, the curve returns to its peak value. These apparent changes in bacterial density are due to aggregation and disaggregation, not an actual change in bacterial cell concentration. Our aggregate correction formula successfully eliminates these spurious abundance changes.

In Fig. S4 and Fig. S5 we reproduce Figures 2 & 4 from the main text to show that the primary results of this work do not depend on the application of the aggregate correction algorithm. The only case in which there is qualitative disagreement between aggregate-corrected B abundance and not-aggregate-corrected B abundance is the 1600 Lux (low light) $t_{inv}$=4d invasion of bacteria on AC as depicted in panel (c) of both Fig. 2 (main text) and Fig. S4. In the aggregate-corrected case, Fig. 2c (main text), B invades and rises to an abundance of $\sim 7 \times 10^5\,\mathrm{mL}^{-1}$. In contrast, in the not-aggregate-corrected case, Fig. S4c, B does not rise significantly above its abundance at introduction and only reaches $\sim 1 \times 10^5\,\mathrm{mL}^{-1}$. The two possible explanations for this disagreement are (1) the B in this case are highly

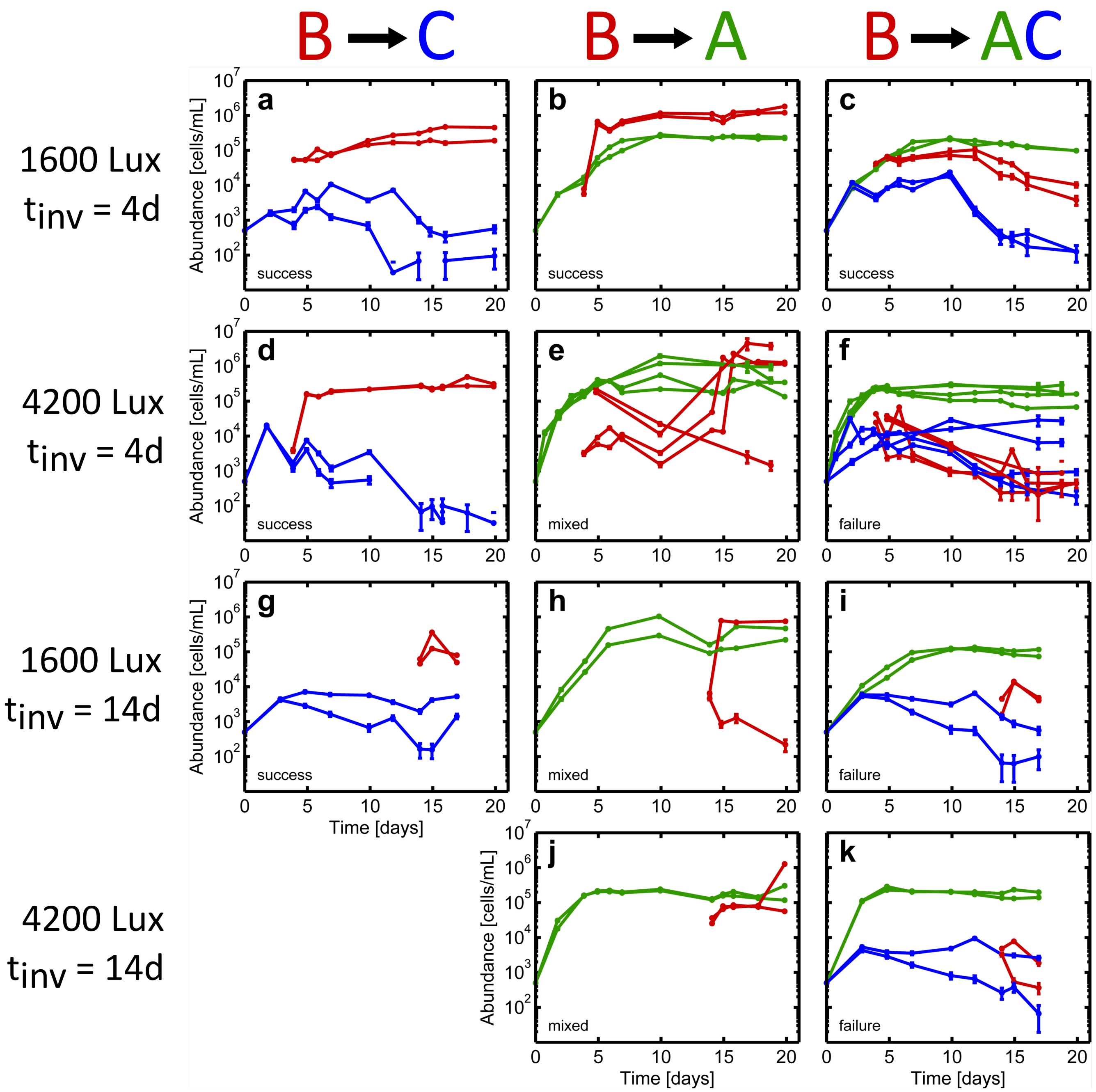


FIG. S4: **Figure 2 from the main text but without the aggregate correction algorithm applied** Panels are identical to Figure 2 of the main text.

aggregated and the aggregate correction algorithm has successfully estimated the true B abundance or (2) high SSC detritus from A and/or C has bled into the flow cytometry gate used to count B and has erroneously inflated the measure of B abundance by $\sim 6 \times 10^5\,\mathrm{mL}^{-1}$.

In order to test explanation (2), we applied the flow cytometry gate used to count B to 1600 Lux (low light) AC coculture data. At almost all timepoints in this data, no detritus fell within this gate. In just two cases did any AC detritus bleed into this gate and at most, $3 \times 10^3\,\mathrm{mL}^{-1}$ B abundance (aggregate-corrected) was erroneously measured. This value is two orders of magnitude smaller than necessary to explain the mismatch between Fig. 2c (main text) and Fig. S4c and so we therefore conclude that the successful invasion depicted in Fig. 2c (main text) is real.

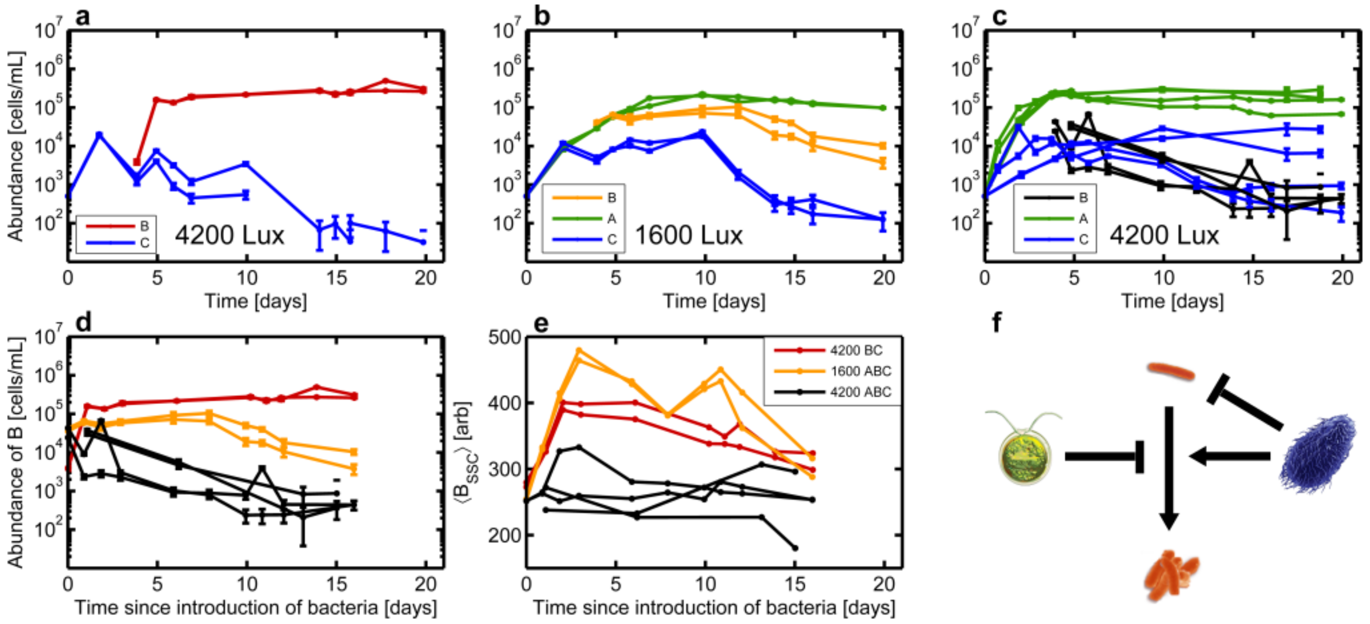


FIG. S5: **Figure 4 from the main text but without aggregation correction algorithm applied** Panels are identical to Figure 4 of the main text.

##### Ciliates cause bacteria to aggregate

When bacteria from a 1600 Lux (low light) B monoculture (Fig. S6a) are compared to bacteria from a 1600 Lux B invasion of C (Fig. S6b), histograms of the side-scatter signal of bacteria show increased aggregation in the presence of the ciliates (Fig. S6g). Note that both histograms have a peak at low side scatter signal which corresponds to planktonic bacterial cells.

##### Ciliate induces aggregation in a Δ*csgA* *E. coli* mutant

In order to investigate the seeming necessity of bacterial aggregation in order to survive ciliate predation, we constructed a Δ*csgA* strain (in an MG1655 background). *csgA* encodes a structural subunit of the curli fimbrae which mediate cell-cell adhesion at temperatures below 37 °C (our experiments were all undertaken at 30 °C)[1]. As expected, monocultures of this mutant aggregate much less than monocultures of the strain used in all other experiments ($\Delta flu \Delta fimA$) (Fig. S7e). See the Methods section of the main text for details of strain construction and genotypes.

Because of its lack of aggregation, we hypothesized that this non-aggregating mutant would be unable to invade a culture of ciliates. Surprisingly, we found that not only did the Δ*csgA* bacteria invade the ciliates (Fig. S7d), but they also showed enhance aggregation in the presence of C as compared to monoculture (Fig. S7e&f).

#### LIGHT LEVEL DOES NOT AFFECT BACTERIAL OR CILIATE ABUNDANCE DYNAMICS

Experiments throughout the paper are performed at 1600 Lux (low light) and 4200 Lux (high light). Here we show that the abundance dynamics of B and C are not impacted by illumination at either level of illumination. When comparing bacterial abundance dynamics between low light and high light conditions for the co-culture experiments, B behaves identically across light levels regardless of the species composition: B monoculture, AB co-culture, or BC co-culture (Fig. S8a,b&c). The only species composition in which B behaves differently across light levels is the ABC co-culture (Fig. S8d). We argue in the main text that this effect is due to a higher-order interaction in the community.

The same independence of dynamics relative to light level is true for ciliates (Fig. S8). We note that there is substantial variability in the C abundance dynamics. We believe that these differences reflect differences in the state

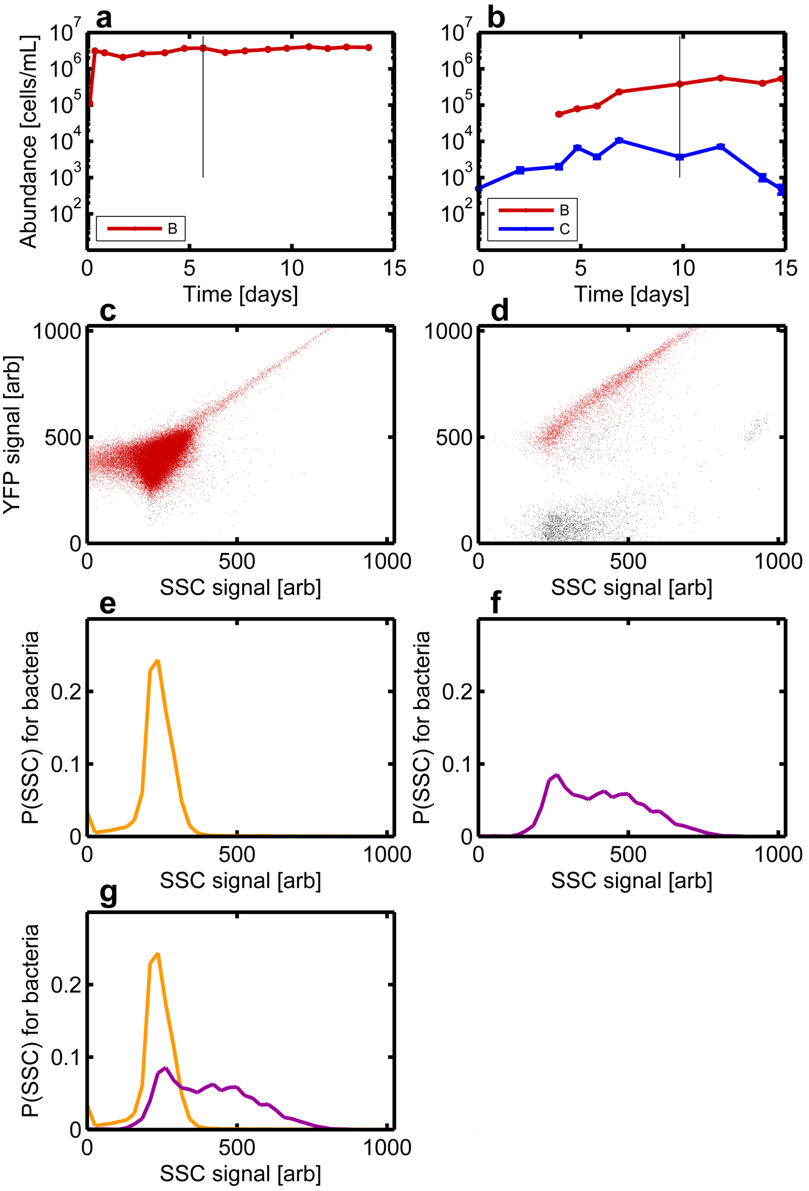


FIG. S6: **Ciliates induce bacteria to aggregate a,** Abundance dynamics for a 1600 Lux (low light) monoculture of B. Black line indicates the time point for which we plot flow cytometry data in **c.** **b,** Abundance dynamics for a 1600 Lux (low light) B invasion of C. The black line indicates the time point for which we plot flow cytometry data in **d.** **c,d** Plotting yellow fluorescence (YFP) versus side-scatter (SSC) for flow cytometry data taken from the indicated day in **a** and **b** respectively. Red-colored points indicate objects we have labeled as bacteria. **e,f**, Histograms of the SSC signal for all objects labeled as bacteria in **c** or **d**. **g,** Overlay of the histograms.

of the ciliate population prior to starting the experiment (e.g. small variations in the growth phase at time the experiment was initiated).

#### ALGAE-BACTERIA INTERACTIONS

##### When algae is at sufficiently high density, it stochastically prevents bacterial invasion

To further investigate the inhibition of B by A we performed a set of invasion experiments where B was introduced to A at $t = 0$ days (co-culture), 1 day, 3 days and 4 days all in high light (4200 Lux) conditions. We found that A did not suppress B when A was at densities below $1 \times 10^5$ mL$^{-1}$ at time of bacterial introduction (Fig. S9 a,b),

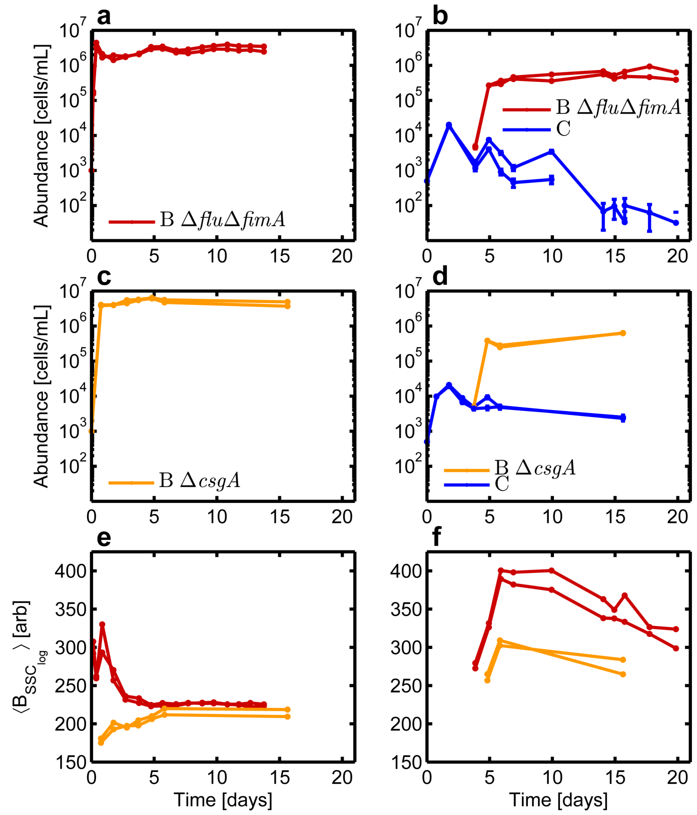


FIG. S7: **Ciliate induces aggregation in Δ*csgA* *E. coli*** All experiments in this figure are at 4200 Lux (high light). **a,** Abundance dynamics for two replicates of a B monoculture, using strain $\Delta flu\Delta fimA$. This is the strain used throughout this study. **b,** Abundance dynamics for two replicates of a B invasion on C, also using the $\Delta flu\Delta fimA$ strain of B. **c,** Abundance dynamics for two replicates of a B monoculture, using strain $\Delta csgA$. **d,** Abundance dynamics for two replicates of a B invasion on C, also using the $\Delta csgA$ strain of B. **e,** Mean side-scatter plotted versus time for the bacteria in panels **a** and **c**. **f,** Mean side-scatter plotted versus time for the bacteria in panels **b** and **d**.

but that suppression did occur when B was introduced to high density A (Fig. S9 c,d). Of those six high-density A cultures which suppressed bacterial invasion, three completely prevented bacterial invasion (low bacterial densities of $\sim 1 \times 10^3\,\mathrm{mL}^{-1}$ even after two weeks), while the other three high-density A cultures allowed bacteria to grow to high density over the period of approximately two weeks following bacterial introduction.

##### Algae must be physically present and illuminated to inhibit bacterial invasion

We performed an experiment to test the importance of the physical presence of algae in the suppression of bacterial invasions. A culture of algae was grown in a 1 L Erlenmeyer flask for ten days in a shaker incubator at approximately 4000 Lux, 30 °C, and 175 RPM. The algae culture was then transferred to vials and inoculated with bacteria at a density of $1 \times 10^5\,\mathrm{mL}^{-1}$. These vials were placed in the culture devices used for the experiments shown in the main text. In two replicates the brightness was set to 4200 Lux (high light) (Fig. S10a) while in the other two replicates the brightness was set to 0 Lux (no light) (Fig. S10b). Bacteria and algae abundance were then measured by flow cytometry.

In the other half of the experiment, the algae culture was filtered through a 0.22$\mu$m PES membrane filter before being distributed across vials and inoculating bacteria. Once again the vials were set to 4200 Lux (Fig. S10c) and 0 Lux (Fig. S10d).

The only condition in which bacterial invasion was inhibited was the condition with lights on and algae physically present (Fig. S10a). In all other cases: lights-off, algae filtered out, or both, bacteria invaded immediately. The necessity of light implies that algae's photosynthetic metabolism must be active to suppress bacterial invasion.

The mechanism by which the physical presence of algae is necessary to inhibit invasion remains unclear. However,

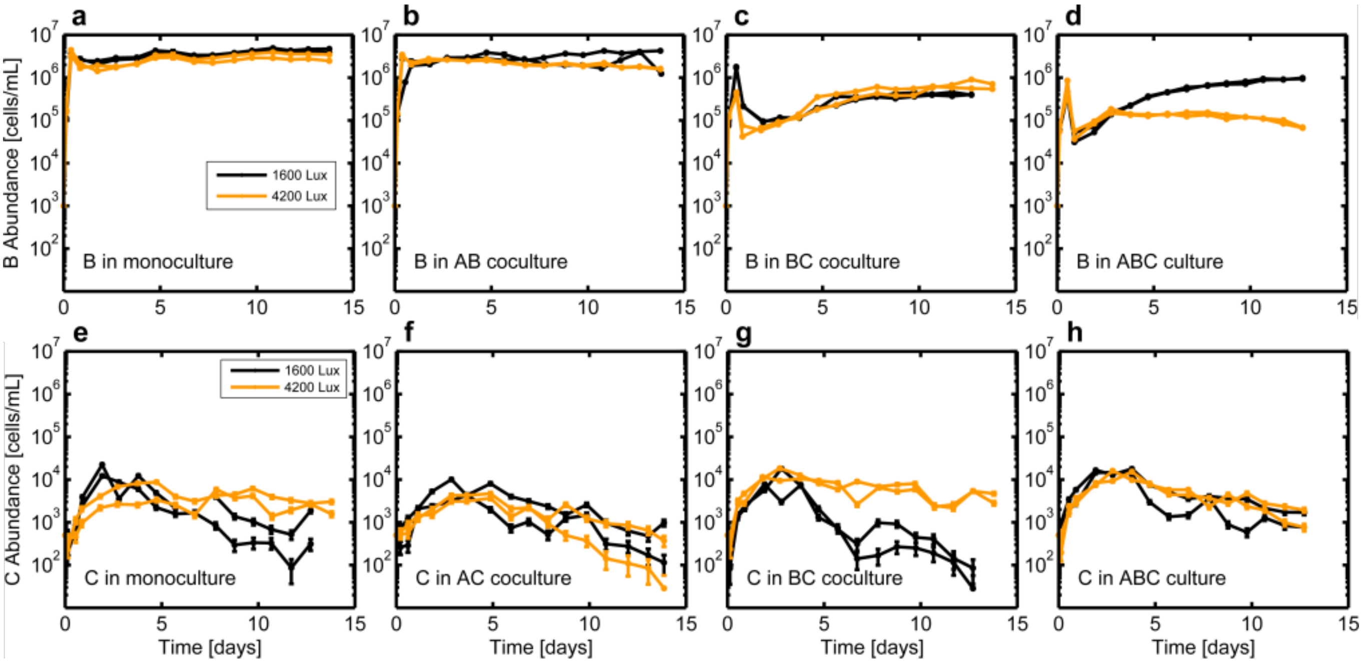


FIG. S8: **Bacterial or ciliate abundance dynamics with different light levels and community composition a-d,** Abundance dynamics for two replicates each of B in 1600 Lux and in 4200 Lux B monoculture (**a**), AB co-culture (**b**), BC co-culture (**c**), and ABC culture (**d**). Legend in **a** applies to **b-d**. **e-h,** Abundance dynamics for two replicates each of C in 1600 Lux and in 4200 Lux C monoculture (**e**), AC co-culture (**f**), BC co-culture (**g**), and ABC co-culture (**h**). Legend in **e** applies to **f-h**.

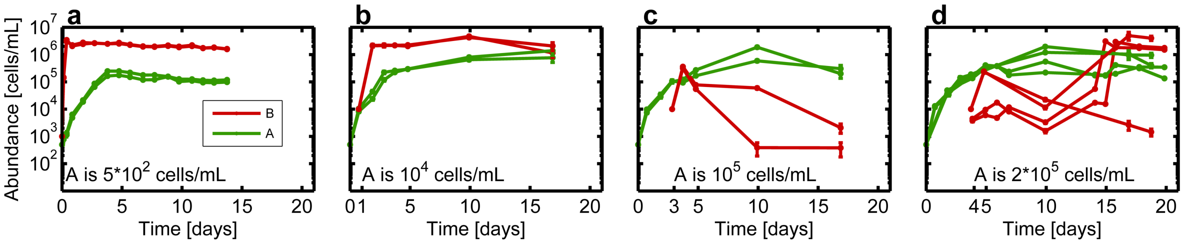


FIG. S9: **Outcome of bacterial invasion in high density algal cultures is stochastic** Abundance dynamics of algae and bacteria when bacteria is grown in co-culture with algae (**a**), or introduced into an algae monoculture at day 1 (**b**), day 3 (**c**), or day 4 (**d**). Text inside panels indicates density of algae at time of introduction of bacteria. All experiments in this figure performed at 4200 Lux (high light).

we present a few possible interpretations of this result: (1) Algae suppresses bacterial invasion by secreting a toxic compound, one that they only begin producing when they sense the presence of bacteria (microbes can be stimulated to emit toxins by presence of other microbes[2]) (2) Algae suppresses bacterial invasion by secreting a toxic compound which can be degraded by bacteria. In this scenario the toxin in the spent algal medium is rapidly degraded by the bacteria, but in the case where algae are present the production rate of the toxin exceeds the bacterial degradation rate of that toxin. This possibility motivated us to test $H_2O_2$ as the mechanism since algae produce reactive oxygen species and bacteria degrade them via catalase. Our experiments showed that hydrogen peroxide is not the mechanism of inhibition (see discussion below and Fig. S12). (3) Physical contact between algae and bacteria is necessary for the mechanism of invasion suppression (flagella have been seen to mediate interactions between microbes[3]). (4) The most pathological possibility: algae suppresses bacterial invasion by secreting a toxic compound that is larger than the pores of the 0.22$\mu$m filter.

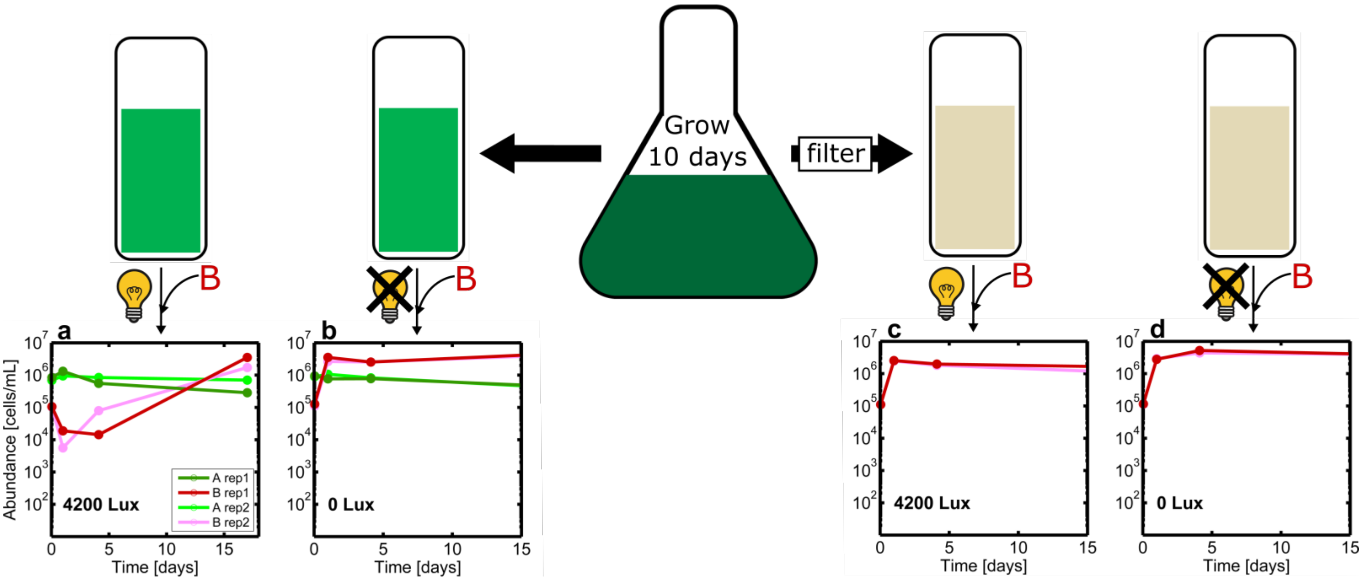


FIG. S10: **Algae must be physically present and illuminated to inhibit bacterial invasion** Algae are grown for 10 days in large volume in a flask of 1/2xTaub .01%pp3 at 30 °C in a shaker-incubator while being illuminated at approximately 4000 Lux. Half the culture is filtered. Then unfiltered (**a,b**) and filtered (**c,d**) algae culture is distributed across vials. Bacteria is inoculated at density $1 \times 10^5$ $\mathrm{mL}^{-1}$ and the vials are then grown at 4200 Lux (high light) (**a,c**) or 0 Lux (no light) (**b,d**). Abundance dynamics after inoculation with bacteria are plotted. There are two replicates for each condition.

##### There exists a threshold light level below which algae cannot suppress bacterial invasion

We performed an experiment to test the importance of light when it comes to algae suppressing bacterial invasion. A culture of algae was grown in a flask (1 L) for ten days in a shaker incubator at approximately 4000 Lux, 30 °C, and 175 RPM. The algae culture was then transferred to vials and inoculated with bacteria at bacterial density $2 \times 10^4\,\mathrm{mL}^{-1}$. These vials were placed in the temperature/light-control systems that the invasion and co-culture experiments in the main text were performed in. In sets of two replicates the brightness was set to 600 Lux (extra low light) (Fig. S11a), 1600 Lux (low light) (Fig. S11b), 2900 Lux (medium light) (Fig. S11c), and 4200 Lux (high light) (Fig. S11d). Bacteria and algae abundance were then measured by flow cytometry.

Algae was only able to suppress bacterial invasion at the highest light level (Fig. S11d). At all other light levels the bacteria invaded immediately. This contradicts the results of the main text in the sense that in experiments reported in the main text, 1600 Lux monocultures of algae were able to suppress bacterial invasion as long as the algae density was high enough, whereas here, for example (Fig. S11b), a 1600 Lux monoculture of high-density algae was not able to suppress bacterial invasion. This result implies that the physiology of algae grown in the flask in the shaker-incubator is significantly different from the physiology of algae grown in the temperature/light-control systems and that the suppression of bacterial growth by algae depend on the growth history of the algal culture or the precise culture conditions.

##### Hydrogen peroxide is not responsible for A inhibiting B invasion

Because metabolically active algae are known to produce reactive oxygen species[4], we tested if $H_2O_2$ was responsible for algae's ability to suppress bacterial invasion. To measure $H_2O_2$ we used the iodine based absorbance method of Junglee et. al.[5]. Absorbance values from cultures were compared to those taken for solutions with known concentrations of hydrogen peroxide.

In a control experiment we determined the concentration of $H_2O_2$ necessary to inhibit bacterial growth. Bacteria were inoculated into a 96-well plate in wells that contained 1/2x Taub .01% proteose peptone No. 3 and $H_2O_2$ in concentrations that ranged from 1M to 1nM. We inoculated bacteria at both $1 \times 10^4\,\mathrm{mL}^{-1}$ and $1 \times 10^5\,\mathrm{mL}^{-1}$. The bacteria were grown in a plate reader at 30 °C and abundance was measured continuously via absorbance at 600 nm. For the low B inoculum, 1mM initial $H_2O_2$ concentration was necessary to prevent growth while for the high B

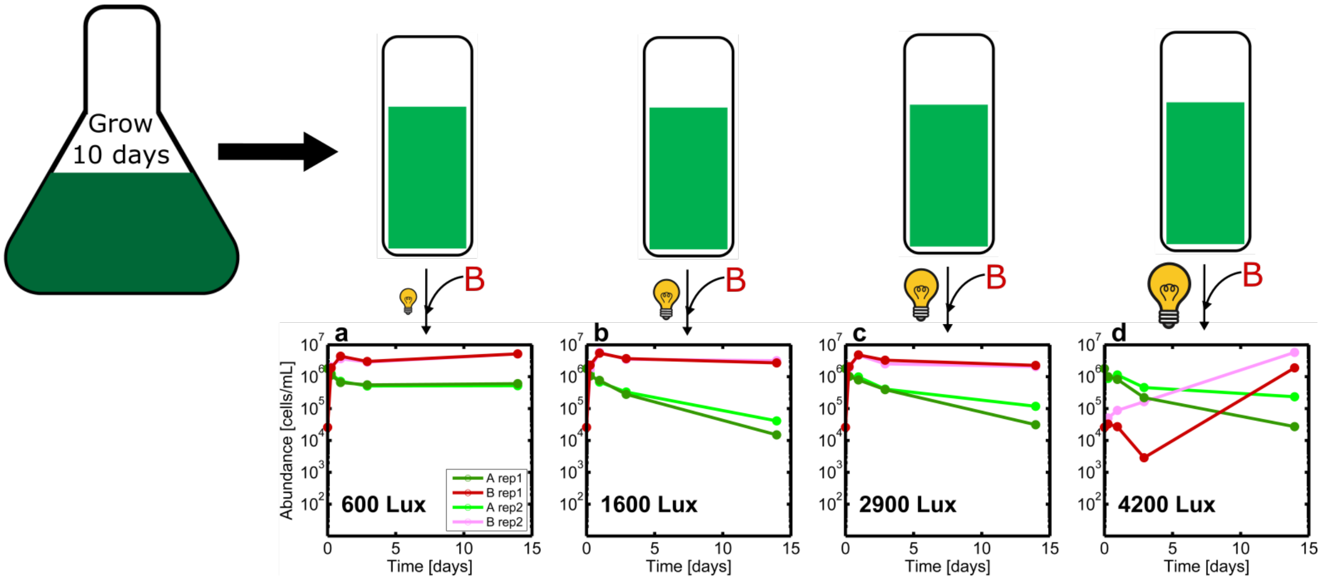


FIG. S11: **There exists a threshold light level below which algae cannot suppress bacterial invasion** Algae are grown for 10 days in large volume in a flask of 1/2xTaub .01% proteose peptone No.3 at 30 °C in a shaker incubator while being illuminated at approximately 4000 Lux. Algae culture is then distributed across vials. Bacteria is inoculated at density $2 \times 10^4\,\mathrm{mL}^{-1}$ and the vials are then grown at 600 Lux (extra low light) (**a**), 1600 Lux (low light) (**b**), 2900 Lux (medium light) (**c**), or 4200 Lux (high light) (**d**). Abundance dynamics after inoculation with bacteria are plotted. There are two replicates for each condition.

inoculum 10mM initial $H_2O_2$ concentration was necessary to prevent growth (Fig. S12a). Measurements of $H_2O_2$ in these cultures one day after inoculation showed that bacteria who successfully grew eliminated the $H_2O_2$ (Fig. S12b).

We then measured the $H_2O_2$ in conditions where B successfully invaded A and also conditions where its invasion was inhibited by A. Specifically, we took $H_2O_2$ measurements in the experiment from Fig. S11. No $H_2O_2$ was detected at any point in any of the systems, and thus we conclude that $H_2O_2$ is not the mechanism by which algae suppresses bacterial invasion. We cannot rule out other reactive oxygen species by this assay.

##### Algal-bacterial adhesion and invasion supression

By closely examining flow cytometry data from the experiment in Fig. S11, it can be seen that bacteria are sticking to algae. Recall that in this experiment algae were grown in a flask for ten days and were then distributed across vials. These vials were inoculated with bacteria and then placed in the temperature/light-control systems at four different light levels. By looking at flow cytometry data taken 6 hours into the experiment, one can see that some of the bacteria have stuck to algae. This is evident from the presence of a small cloud of objects distinct from algal signals that are both high YFP and chlorophyll. At this time-point, the bacteria form a tight cloud around YFP = 500 (Fig. S13b,e). When YFP signal is plotted versus Chlorophyll signal for flow cytometry from this same time point, one can see that there is a cloud with that same YFP value, that also has high chlorophyll, and is distinct from the main cloud of algae points (beige points, Fig. S13c,f). These facts taken together indicate a fraction of the bacteria have stuck to the algae. The reader may wonder why, when both bacteria and algae have significant YFP signal, the cloud of B-stuck-to-A objects does not appear on the plot to have a YFP value that is the sum of both. Recall from the aggregate correction section of this supplement that flow cytometry signals for an object should be understood as the log of an object's intensity for a given scattering/fluorescence channel, and thus values do not add linearly on these plots.

We next asked: does the degree of bacteria sticking to algae depend on the degree to which the invasion is inhibited? Could differential sticking be responsible for invasion suppression? In the experiment from Fig. S11, there were six immediately successful invasions, one of which is depicted in Fig. S13a, and two suppressed invasions, one of which is depicted in Fig. S13d. We used a strict gate to determine the size of the population of bacteria stuck to algae (Fig. S13c&f). For the six systems shown in Fig. S11 where bacteria invade successfully immediately, we find that the fractional abundances of bacteria stuck to algae ($\frac{B-stuck-to-algae}{B-not-stuck-to-algae}$) are: 3.7 ± 0.1%, 3.7 ± 0.1%, 2.8

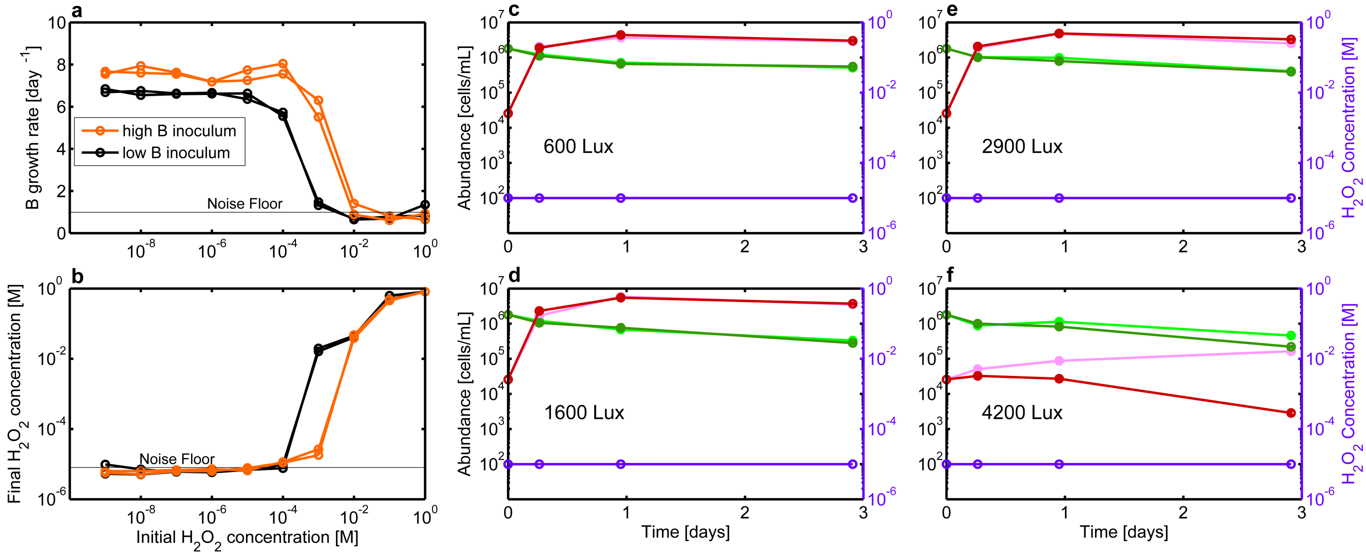


FIG. S12: **$H_2O_2$ is not the mechanism through which A inhibits B growth a,** Growth rate of bacteria as a function of the initial $H_2O_2$ concentration for both high and low inoculum. Growth media is 1/2x Taub .01% proteose peptone No. 3. High B inoculum is $1 \times 10^5\,\mathrm{mL}^{-1}$, while low B inoculum is $1 \times 10^4\,\mathrm{mL}^{-1}$. Noise floor in growth rate was calculated by measuring the growth rate (via regression) on synthetic data generated from a normal distribution with the same mean and variance as observed experimentally (e.g. absorbance fluctuations in wells where no growth occurs). **b,** Final $H_2O_2$ concentration as function of the initial $H_2O_2$ concentration in the same experiment as **a**. When final $H_2O_2$ concentrations are calculated based on absorbance measurements, background noise in the absorbance always leads to at least $1 \times 10^{-5}$M concentration of $H_2O_2$ being calculated and so that is where we set our noise floor. **c-d** Abundance curves from the experiment in Fig. S11. At each time point, a sample was taken and its $H_2O_2$ level was measured. $H_2O_2$ levels were below our limit of detection at all time points.

± 0.1%, 2.5 ± 0.1%, and 2.9 ± 0.1%. Error bars are estimated assuming Poisson counting error. For the two systems where the invasions were suppressed, we find 4.8 ± 0.9% and 7.7 ± 1.4%. A two-sample t-test assuming unequal variances fails to reject the null hypothesis that the average fraction of B cells stuck to A differs between successful and inhibited invasions ($p = 0.28$). We note the small samples size means this result should not be taken too seriously. However, based on the inconsistency between the two values in the case of the suppressed invasions, and the near-overlap between the error bars of one of the values for suppressed invasion (4.8 ± 0.9%), and two of the values for immediately successful invasion (both 3.7 ± 0.1%), we suggest it is not reasonable to believe that there is a substantially larger fraction of B adhered to A when B invasions are supressed.

#### Physical collisions between bacteria and algae are frequent even when algal density is low

In the main text we conjectured that the density dependence of algal inhibition of bacteria might arise from more frequent cell-to-cell contact when algal densities are high. Here we estimate the frequency of this contact and find that bacteria come in contact with algae with high frequency even at low algal densities.

We calculate the number of algae that a bacterium encounters in one second. From Stocker et. al, $E_B = 4\pi N_A(D_A + D_B)(r_A + r_B)T$ where $E_B$ is the number of algae a single bacterium encounters in time $T$, $N_A$ is the concentration of algae, $D_A$ and $D_B$ are the diffusivity of algae and bacteria respectively, and $r_A$ and $r_B$ are the radii of algae and bacteria respectively. Taking $T$ to be 1 second, we will attempt to calculate a lower bound on $E_B$, the number of algae a bacterium encounters in one second. For $N_A$, we use $500\,\mathrm{mL}^{-1}$ ($5 \times 10^8\,\mathrm{m}^{-3}$), the starting concentration of algae, and therefore the lower bound. From Stocker, $D_B = \frac{U^2\tau}{3}$ where $U$ is the speed of the bacterium and $\tau$ is the turning rate. We take $\tau$ to be one turn per second[6]. In our systems, $U$ is not the swimming speed, but rather the speed imparted on the bacterium through stirring. The stir-bar in the vial turns at 450 RPM and is of diameter 1.5 cm. Assuming that the average bacterium will move at the same speed as the point halfway along the radius of the stir-bar, we obtain speed $0.17\,\mathrm{m\,s}^{-1}$. This value is consistent with the observation that a drop of food coloring applied to the top of the vial while the vial is being stirred distributes throughout the vial instantaneously by eye. $D_B$ is therefore $0.0096\,\frac{\mathrm{m}^2}{\mathrm{s}}$. We set $D_A = 0$ in the interest of establishing a lower bound. We take $r_B$ to be $1\,\mu\mathrm{m}$ and

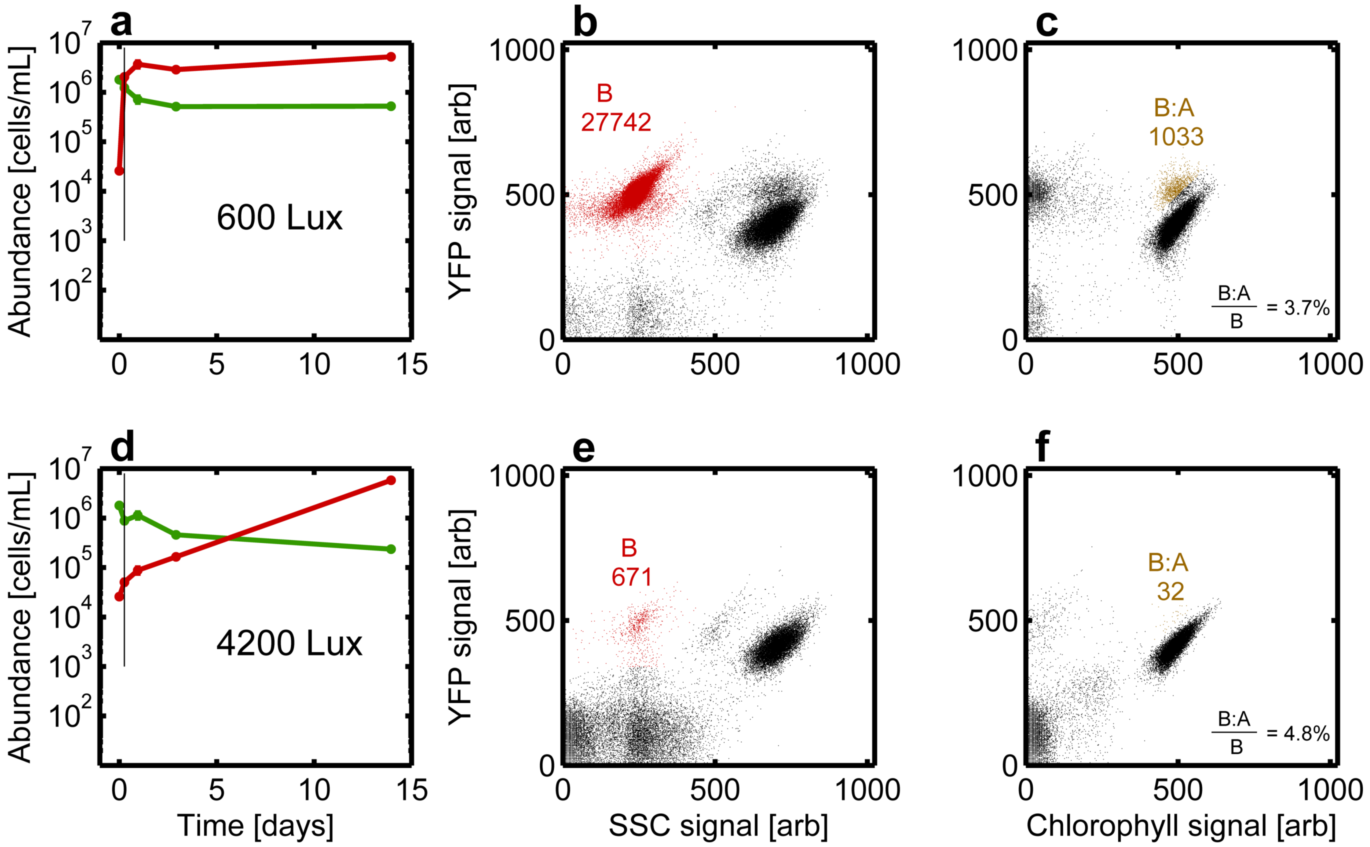


FIG. S13: **A small fraction of bacteria adhere to algae a,** Abundance curves for algae and bacteria taken from one of the 600 Lux replicates of the algae flask experiment in Fig. S11. The black line marks the time point for which flow cytometry data is plotted in **b&c. b,** YFP signal plotted versus SSC signal for flow cytometry data from aforementioned timepoint. Points in red mark objects classified as bacteria. Number indicates the number of these objects. **c,** YFP signal plotted versus Chlorophyll signal for flow cytometry data from that same timepoint. Points in gold mark objects classified as algae with bacteria stuck to them. Number indicates the number of these objects. Percentage indicates how many of these algae-stuck-to-bacteria objects there are as a fraction of the normal bacteria. **d,e&f,** The same analysis as in the top row of this figure, but instead with one of the 4200 Lux replicates from Fig. S11.

$r_A$ to be 1 µm. Taken together, we calculate $\frac{E_B}{T}$ to be 180 collisions per second. That is 180 algae encountered per second by a single bacterium.

#### SPENT MEDIA EXPERIMENTS

##### Neither algae nor ciliates compete with bacteria for nutrients

We considered the possibility that nutrient competition could account for the inhibition of B by A and the failure of B to invade AC communities. In the undefined medium used here it is possible that A and C alone do not consume all of the available carbon and nitrogen but in co-culture they consume an essential nutrient for bacterial growth. To investigate this possibility we performed a series of spent media experiments where communities were grown in our growth chambers, samples were harvested and then all cells were removed by filtration (example shown in Fig. S14a). B was then inoculated into this spent medium and grown to saturation $K_B$ in a 96-well plate where its density was assayed by flow cytometry after two days of growth (Fig. S14a). We find that B is able to grow on the spent medium of A, C and AC communities to a saturating density that is identical to growth on fresh medium irrespective of the time at which the spent media was taken (Fig. S14c). This result rules out the presence of nutrient competition between these three species as the source of the invasion outcomes in the main text. In addition, the fact that B grows rapidly to maximum density in spent medium harvested from A communities at high algal densities shows that nutrient competition is not the cause of A inhibiting B invasions.

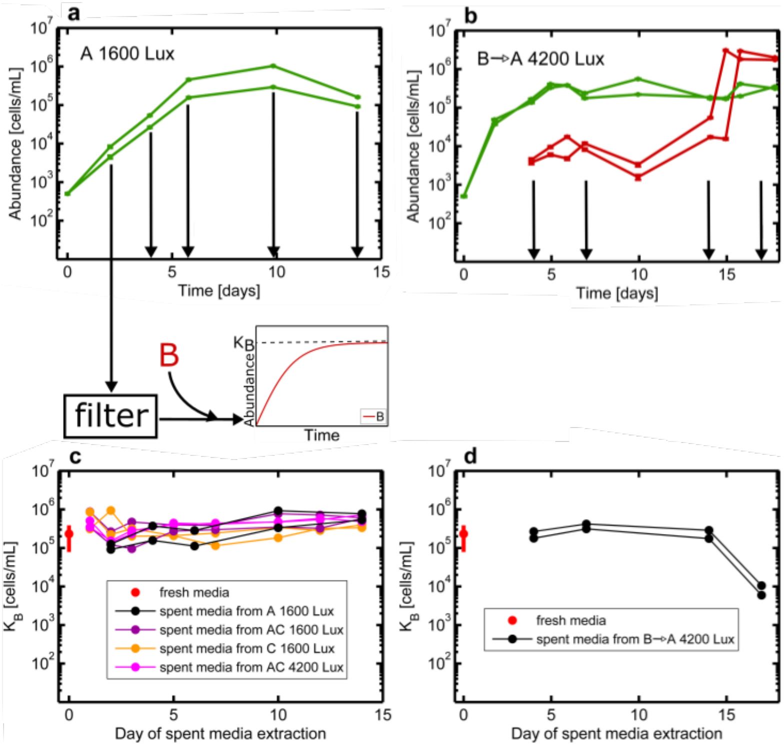


FIG. S14: **Algae and ciliates do not consume bacterial nutrients a,** Abundance plotted versus time for two replicates of an algae monoculture at 1600 Lux (low light). Black arrows indicate the time points when media is extracted and filtered. Bacteria is then grown on this spent media in a microtiter plate until it reaches saturation. The saturating density $K_B$ is measured via flow cytometry. This experiment is also performed on a 1600 Lux C monoculture, a 1600 Lux AC coculture, and a 4200 Lux AC coculture. **c,** $K_B$ is plotted for all the conditions. The x-axis indicates the day of extraction of spent media. **b,** The same spent media extraction experiment is performed on a monoculture of algae that is invaded with bacteria. This means that the spent media taken from this culture has already been exposed to bacteria. **d,** $K_B$ plotted versus day of spent media extraction for the experiment in **b.**

To further investigate the role of nutrients in the inhibition of B by A we harvested spent medium from an A community after a B invasion at several time points (Fig. S14b). We then filtered out both A and B and then inoculated fresh B cells at low density in this spent medium. Remarkably, B was able to grow on spent medium from an AB invasion experiment so long as the spent medium was taken prior to the invading B population reaching high density. After the point where invading B reach high density, B can no longer grow to high density on spent medium (Fig. S14d). This result shows that the inhibition of B by high density populations of A limits the ability of B to consume nutrients. This experiment further suggests that the inhibition of B by A may be caused by a volatile compound or direct physical contact rather than a soluble toxin.

##### Algae spent media de-aggregates bacteria

The spent media experiment shown in Fig. S14a can also be analyzed to show that algae spent media has a de-aggregating effect on B. Two replicates of a 1600 Lux (low light) A monoculture were grown in the temperature/light-control systems. Samples were harvested at multiple time points and all algae cells were removed by filtration. B was then inoculated into this spent media (Fig. S15a) and grown to saturation in a 96-well plate where B's density $K_B$ was assayed by flow cytometry after two days of growth. The mean side-scatter signal of bacteria decreased as a function of time of spent media extraction in both replicates (Fig. S15b). The longer the duration of algal growth the greater is the de-aggregation of B grown on spent medium.

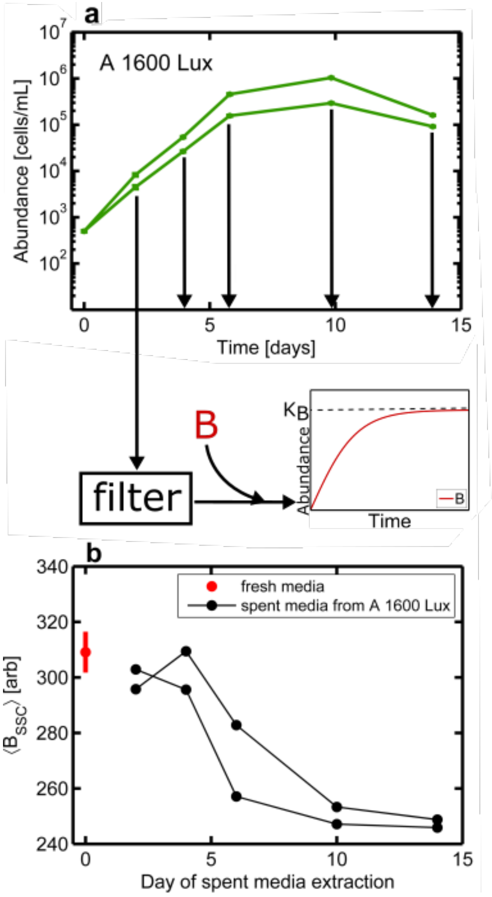


FIG. S15: **Algae spent media de-aggregates bacteria a,** Abundance plotted versus time for two replicates of an algae monoculture at 1600 Lux (low light). Black arrows indicate the time points at which media is extracted and filtered. Bacteria is then grown on this spent media in a microtiter plate until it reaches saturating density $K_B$ **b,** Mean side-scatter of the bacteria after they reach $K_B$ is plotted versus time of spent media extraction. The point at time zero represents the mean-side scatter of bacteria that were grown on fresh media.

##### Ciliates present in algal spent media is not sufficient to prevent bacterial invasion

In the main text we show that a 3-body interaction between algae, bacteria, and ciliates prevents the invasion of the bacteria. We thought to ask if this interaction requires the physical presence of the algae or if spent media from the algae is sufficient. Our objective was to determine whether the impact of algal spent medium de-aggregation altered the predation pressure of B by C.

The experiment was performed in a similar manner to the spent media experiments in the main text. Two replicates of a 1600 Lux (low light) A monoculture were grown in the temperature/light-control systems. Samples were harvested at multiple time points and all algae cells were removed by filtration. B along with C was then inoculated into this spent media (Fig. S16a) and grown to saturation in a 96-well plate where B's density $K_B$ was assayed by flow cytometry after two days of growth. $K_B$ did not vary with the time of spent media extraction from the algae monocultures and $K_B$ in algae spent media was only, on average, 46% lower than for fresh media (Fig. S16b).

#### LIVE-DEAD STAINING EXPERIMENTS

When gating on clouds of points of flow cytometry data, an assumption is made that the cells which are fluorescent are alive and thus a good measure of true abundance of live cells. We base this assumption on the fact that algae cells fluoresce significantly differently in chlorophyll when alive as compared to dead[7]. Nevertheless, we attempt here to use Sytox Green Dye to assay the number of dead cells. Sytox Green is a nucleic acid stain. Sytox Green stains dead cells positively in the YFP channel.

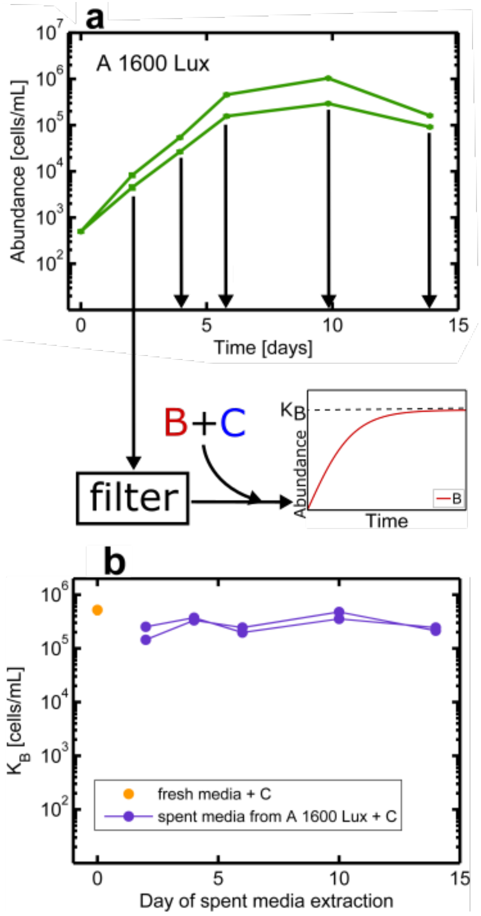


FIG. S16: **Algae spent media alone does not largely alter C predation on B a,** Abundance plotted versus time for two replicates of an algae monoculture at 1600 Lux (low light). Black arrows indicate the time points at which media is extracted and filtered. Bacteria and ciliates are then grown together on this spent media in a microtiter plate, allowing B to reach saturating density $K_B$. **b,** $K_B$, measured by flow cytometry, is plotted versus time of spent media extraction.

We performed the Sytox experiment on algae in four cases, two replicates of a day 4, 1600 Lux (low light) ABC co-culture, and two replicates of a day 4, 4200 Lux (high light) ABC co-culture. In each case, flow cytometry data was taken before the application of Sytox (Fig. S17a), then Sytox Green was added at a concentration of 100nM, incubated for 10 minutes, and then flow cytometry was performed again (Fig. S17b). In each case, a new "dead algae" cloud with saturating YFP signal emerged after application of Sytox. Across the four cases, these dead cells made up 31 $\pm$ 11% of the total algae cells, suggesting up to a third of algal cells are dead. This estimate is likely to be an overestimate due to high stain concentrations as explained below.

The Sytox experiment was also performed on bacteria in four cases, two replicates of a day 14, 1600 Lux B monoculture, and two replicates of a day 14, 4200 Lux B monoculture. Once again a high YFP portion separated out from the main cloud after application of Sytox (Fig. S17c&d). In the case of bacteria, dead cells made up 3.6$\pm$ 2.1% of the total bacterial cells (average and standard deviation across four replicate systems). The Sytox experiment was also performed on ciliates, but the results are impossible to interpret given that Sytox just translated the entire cloud of ciliates to a higher YFP region (Fig. S17e&f) and no separation is observed.

We believe that in the case of algae and ciliates, Sytox concentration was too high, and thus the estimate of dead cells is an overestimate. Note what happens when a 5-fold higher Sytox concentration is used to stain algae (Fig. S18c). The estimate of percentage of dead cells increases from 31 $\pm$ 11% to 51 $\pm$ 21% (average and standard deviation across four replicates). This increase indicates that we are in a regime of high Sytox concentration where even live cells are being stained with Sytox Green. Further supporting this conclusion is the fact that the main cloud of algae points in Fig. S18b&c is also becoming distorted toward high YFP, this arises from excess Sytox staining live cells.

The Sytox Green staining protocol suggests performing repeated staining experiments at varying concentrations of dye and choosing the highest concentration where a subset of the presumably live cells are not stained. Further, the

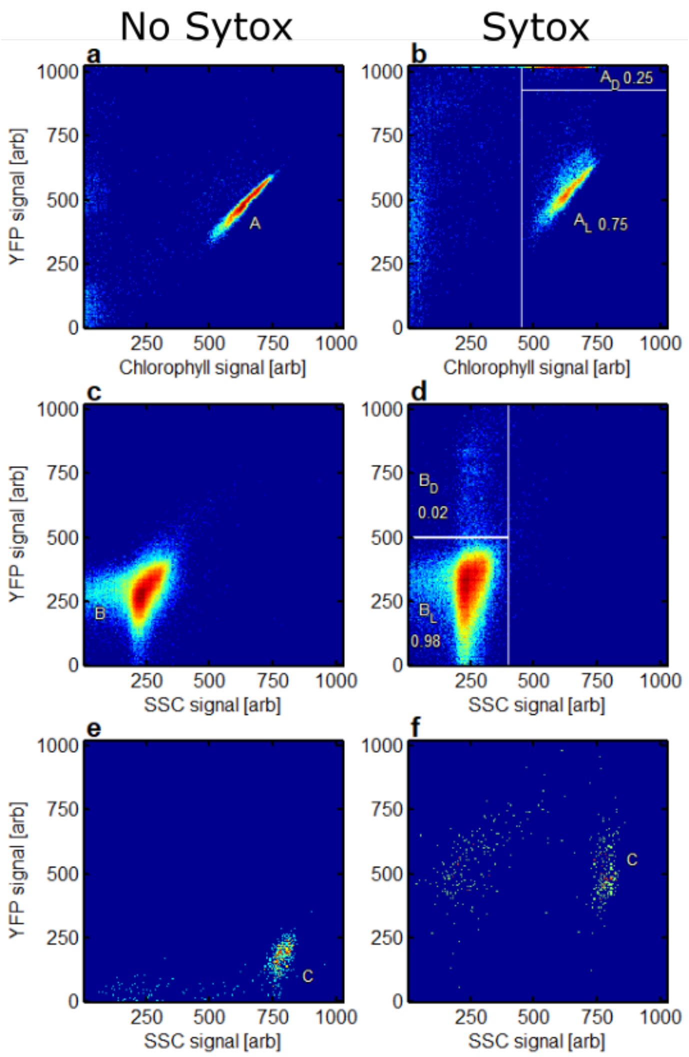


FIG. S17: **Live-dead staining of algae and bacteria a,** YFP signal plotted versus Chlorophyll signal for flow cytometry data taken from day 4 of a 4200 Lux ABC coculture. The cloud labeled $A$ is known to be algae **b,** YFP signal plotted versus Chlorophyll signal for that same sample but after the application of 100 nM Sytox. At sufficiently low concentration, Sytox selectively increases the YFP signal of dead cells. Note how a cloud with saturating YFP signal, thought to be dead algae $A_D$, has separated from the main cloud of live algae $A_L$. Numbers indicate fraction of counts within corresponding white rectangle divided by total counts across both white rectangles. **c,d** A similar analysis is performed on bacteria by plotting YFP signal versus SSC signal. In this case, the sample is taken from day 14 of a 4200 Lux B monoculture and in panel **d** 500 nM Sytox is used. **e,f** A similar analysis is performed on ciliates. In this case, the sample is taken from day 4 of a 1600 Lux C monoculture. Unstained cells are shown in **e** and cells stained with 100 nM Sytox in panel **f**. Note how the entire ciliate cloud shifts in YFP rather than separating out into high YFP and low YFP clouds, thus making quantification of dead cells impossible.

ability of cells to expel the dye depends on their physiological state (e.g. exponential versus stationary phase). Since this state varies throughout our experiment we deemed it infeasible to perform Sytox staining at many time points for the many different conditions studied here. However, the staining data we do have supports the claim for algae and bacteria that less than ∼30 % of the algal population or ∼5 % of the bacterial population are dead. We note that these fractions are small relative the the changes in abundances that occur during our experiment.

#### SUCCESSFUL BACTERIAL INVASIONS ARE NOT AN ARTIFACT OF MIS-CLASSIFING ALGAE DETRITUS AS BACTERIA

Given that detritus accumulates over the course of experiments, we looked closer at flow cytometry data to confirm that accumulating algae detritus was not being mislabeled as a successfully invading bacterial population, especially

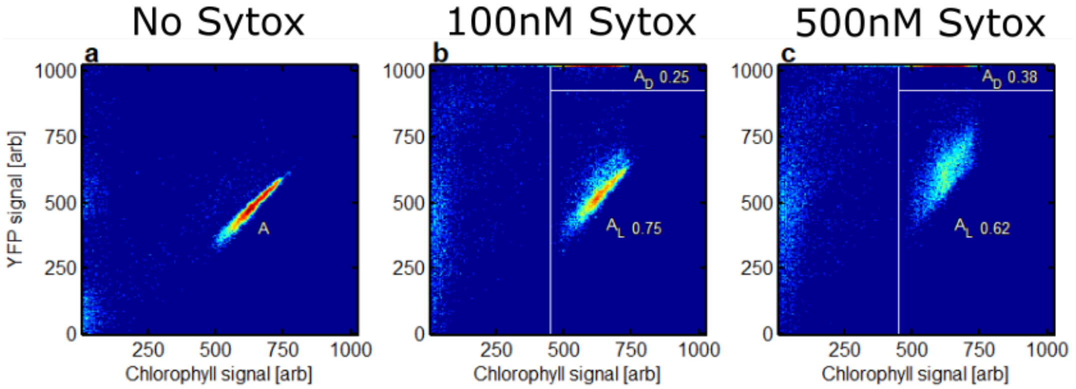


FIG. S18: **Sytox staining overestimates fraction of algal cells that are dead a,** YFP signal plotted versus Chlorophyll signal for flow cytometry data taken from day 4 of a 4200 Lux ABC coculture. The cloud labeled "A" is known to be algae **b,** YFP signal plotted versus Chlorophyll signal for that same sample but after the application of 100 nM Sytox. Note how a cloud with saturating YFP signal comprised of dead algae $A_D$, has separated from the main cloud of live algae $A_L$. Numbers indicate fraction of counts within corresponding white rectangle divided by total counts across both white rectangles. **c,** Same sample and analysis as panel **b** except in this case 500 nM Sytox has been used. Note how the additional Sytox significantly changes the ratio between live and dead cells.

in the case of the stalled bacterial invasion. First we plotted YFP signal versus SSC signal for time points at the beginning, middle, and end of a 1600 Lux (low light) algae monoculture (Fig. S19a-d). We then examined the number of objects that pass a gate for bacteria as a function of time. Over the course of this algae monoculture, only a small number of objects (<100) passed this gate. The same was true for a 4200 Lux (high light) algae monoculture (Fig. S19e-h). In contrast, there were approximately a thousand times as many objects in this gate at the end of a 1600 Lux B invasion on A (Fig. S19i-l) and a 4200 Lux B invasion on A (Fig. S19m-p). These results suggest that algal detritus is not a strong contributor to the number of counts of B we observe by flow cytometry.

#### DETAILS OF MODEL OF ABC DYNAMICS WITH HIGHER ORDER INTERACTIONS

We present a simple, deterministic, ODE based model of population dynamics in the ABC community that reproduces the core features of our data. These features are as follows:

- Bacteria successfully invade AC communities with low algal densities
- Bacteria fail to invade AC communities when A is at high density
- Bacteria invade A and C monocultures with slower invasions of A monocultures at high A densities
- C induces B aggregation
- A inhibits B aggregation
- There is no competition for nutrients in the system that is responsible for these observations

In order to capture these features with a minimum of freely varying parameters, we constructed the following model in which most parameters can be estimated directly from data.

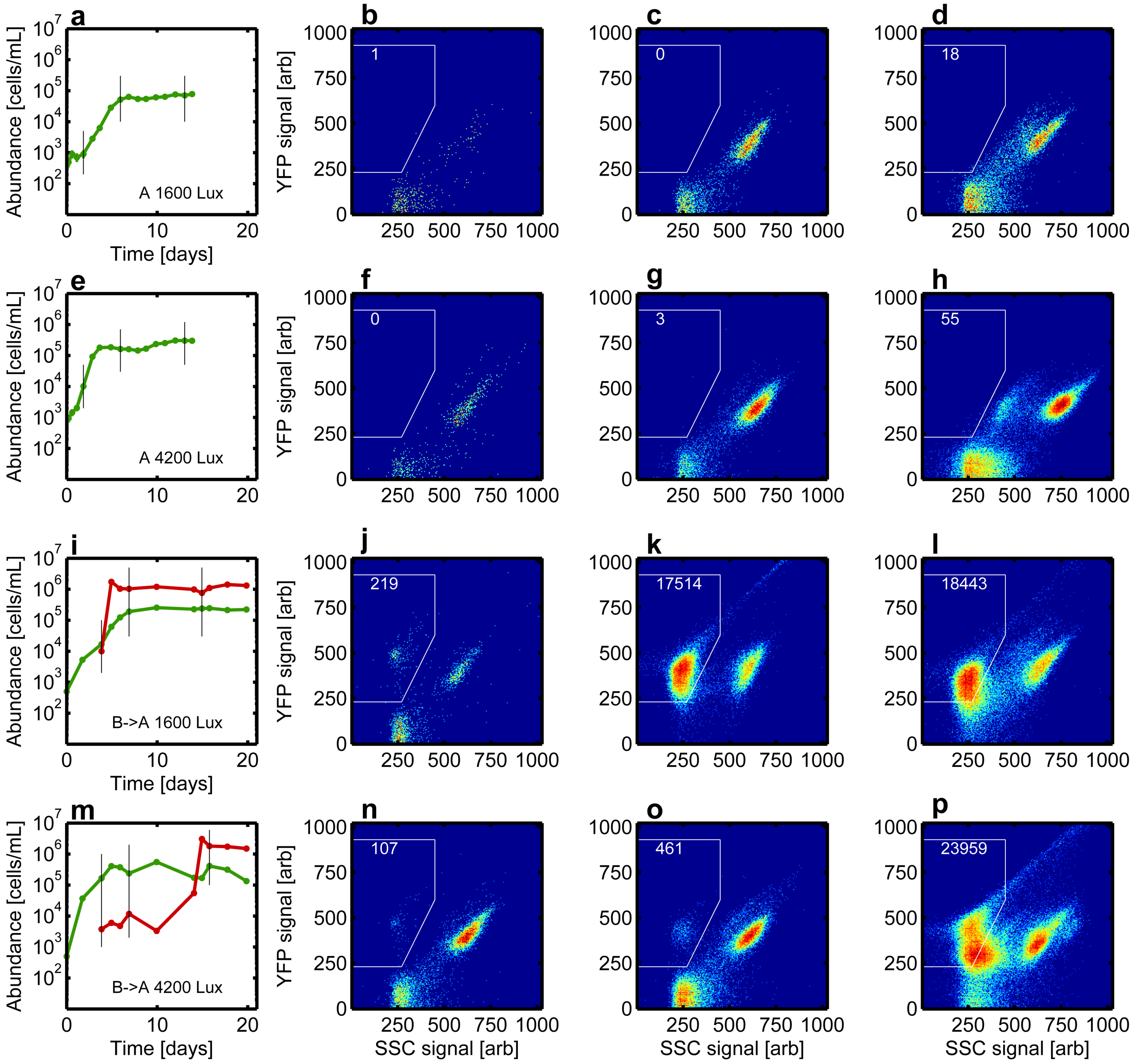


FIG. S19: **Accumulating algae detritus is not being misclassified as bacteria a,** Abundance dynamics for a 1600 Lux (low light) algae monoculture. The three black lines mark the three time points for which flow cytometry data is plotted in **b,c,&d**. **b,** Heatmap for flow cytometry data taken from first marked time point with YFP signal plotted versus SSC signal. Number represents number of counts inside white polygon which is the gate used to detect bacteria. **c,** Same plot for second time point. **d,** Same plot for third time point. Subsequent rows in this figure depict the same analysis, but for a 4200 Lux (high light) algae monoculture, a 1600 Lux B invasion on A, and a 4200 Lux B invasion on A. In all panels with flow cytometry data, the same gate is used.

$$\dot{x}_B = x_B(r_B - r_{AB}\frac{x_A}{K_A})S - Fx_Bx_C - \alpha_1 x_B x_C + \alpha_2 A_B x_A \tag{1}$$

$$\dot{A}_B = \alpha_1 x_B x_C - \alpha_2 A_B x_A \tag{2}$$

$$\dot{x}_A = r_A x_A(1 - \frac{x_A}{K_A}) \tag{3}$$

$$\dot{x}_C = r_C x_C(1 - \frac{x_C}{K_C}) \tag{4}$$

$$\dot{S} = -\frac{x_B}{Y}(r_B - r_{AB}\frac{x_A}{K_A})S \tag{5}$$

$$\tag{6}$$

$x_B$ is the density of planktonic (single-celled) bacteria, while $A_B$ is the density of bacteria in aggregates. $x_A$ and $x_C$ are the density of algae and ciliates respectively. Note that the substrate which drives bacterial growth $S$ is assumed unitless without loss of generality with an initial value of 1. $Y$ then is the carrying capacity of bacteria in the medium used here ($3.9 \times 10^6\,\mathrm{mL}^{-1}$). Only planktonic bacterial cells ($x_B$) can grow on the substrate $S$. We assume that aggregated bacterial cells ($A_B$) do not grow since *E. coli* biofilms are known to exhibit a physiological state similar to stationary phase[8]. Note that the model makes no claim on how many aggregates there are, nor how many bacteria make up a given aggregate; $A_B$ simply denotes how many cells of bacteria are in an aggregated state. We then obtain the total number of bacterial cells $T_B = x_B + A_B$. $T_B$ is what is plotted for bacterial abundances in all figures in the main text and is the output of our aggregate correction algorithm discussed above. Below we justify the functional forms used in this model and the parameter values we chose for numerical simulation.

The parameters of the model are described in the following table with their corresponding values. We are able to directly measure or use previous work to constrain all parameters except $r_{AB}$, $\alpha_1$ and $\alpha_2$. $r_{AB}$ must be on the same order as $r_B$ in order to observe substantial inhibition of B growth by A so we set this parameter accordingly. $\alpha_1$ is inferred indirectly from the data. We treat $\alpha_2$ as the only free parameter in the model and study our model behavior over a range of values (see below).

Values for all parameters in our simulation are given in Table I.

TABLE I: Model parameter values

| Parameter | Value | Source |
|---|---|---|
| $r_B$ | $0.3\,\mathrm{h}^{-1}$ | This study |
| $r_{AB}$ | $0.29\,\mathrm{h}^{-1}$ | Inferred |
| $r_C$ | $0.073\,\mathrm{h}^{-1}$ | This study |
| $r_A$ (high light) | $0.073\,\mathrm{h}^{-1}$ | Fig. S1 |
| $r_A$ (low light, w/BC) | $0.016\,\mathrm{h}^{-1}$ | Fig. S1 |
| $r_A$ (low light, w/C) | $0.025\,\mathrm{h}^{-1}$ | Fig. S1 |
| $r_A$ (low light, w/B) | $0.031\,\mathrm{h}^{-1}$ | Fig. S1 |
| $r_A$ (low light, alone) | $0.045\,\mathrm{h}^{-1}$ | Fig. S1 |
| $K_A$ | $2.3 \times 10^5\mathrm{mL}^{-1}$ | This study |
| $K_C$ | $1.2 \times 10^4\mathrm{mL}^{-1}$ | This study |
| $Y$ | $3.9 \times 10^6\mathrm{mL}^{-1}$ | This study |
| $F$ | $1 \times 10^{-5}$ mL h$^{-1}$ | [9][10] [11], Inferred |
| $\alpha_1$ | $2.5 \times 10^{-6}$ mL h$^{-1}$ | This study |
| $\alpha_2$ | $2 \times 10^{-8}$ mL h$^{-1}$ | Free parameter |

**Bacteria-ciliate interactions**

To begin, we ignore the algae and examine the interaction between the bacteria and the ciliates. This interaction is characterized by five parameters: $r_B$, $Y$, $\alpha_1$, $F$, $r_C$ and $K_C$. Of these parameters, $r_B$, $Y$, $r_C$, $K_C$ and $F$ can be inferred from data acquired for this study. For a more detailed discussion of the estimate of $F$, the feeding rate, from our data, see below. This leaves $\alpha_1$, the rate at which the ciliates induce $B$ aggregation, unknown. We therefore performed simulations on a co-culture of B and C while varying the parameter $\alpha_1$ in order to see which value of

$\alpha_1$ best reproduces the data. The results of these simulation are shown in Fig. S20. The simulations show an intuitive result. In all cases, the bacteria ($T_B$) grow to a high density before the ciliates ($x_C$) have grown to an appreciable density. The aggregation rate $\alpha_1$ then determines how much the bacterial density crashes after that peak. If the aggregation rate $\alpha_1$ is low, the bacteria fail to aggregate sufficiently quickly to avoid predation and the total bacterial density $T_B$ crashes severely. Conversely, if $\alpha_1$ is set high, bacteria aggregate quickly and avoid predation and experience only a very mild fall after the abundance peak. Typically in data when B is grown in coculture with C, we observe bacterial densities peaking to just above $1 \times 10^6\,\mathrm{mL}^{-1}$ before dropping to about $1 \times 10^5\,\mathrm{mL}^{-1}$ (Fig. S8c). Matching simulations to these values is one criteria for choosing $\alpha_1$. The other criteria is as follows. In BC co-culture experiments we observe a transient spike (i.e. the peak followed by crash) in bacterial density while in invasion experiments of B on C the bacterial density does not exhibit a spike (e.g. Fig. 2 , main text). The absence of the spike in bacterial abundances in invasion experiments is likely due to increased initial bacterial aggregation in invasion experiments which is itself driven by the fact that the ciliates are at higher density in invasion experiments at the time of bacterial introduction. Taking both criteria into account, we want the aggregation rate $\alpha_1$ to be low enough that bacterial density drops to $1 \times 10^5$ mL$^{-1}$ in BC co-culture simulations, but high enough that the bacteria do not exhibit a spike in B-invasion-of-C simulations (Fig. S20 bottom right panel). We take $\alpha_1 = 2.5 \times 10^{-6}$ mL h$^{-1}$ as the best compromise between these two criteria and fix this parameter at that value for all future simulations.

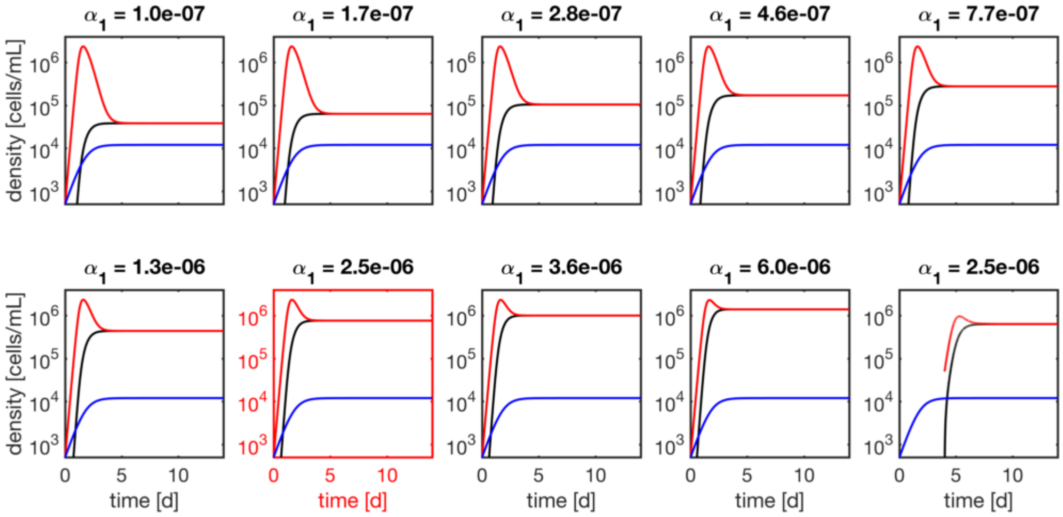


FIG. S20: Abundance dynamics of B and C in co-culture simulated using the model discussed above. Parameter values are as shown in Table I with the exception of $\alpha_1$ which is varied as indicated in the title of each panel (units are mL h$^{-1}$). In each panel, the total bacteria $T_B$ is in red, the ciliates $x_C$ in blue, and aggregating bacteria $A_B$ in black. The bottom right panel shows abundance dynamics of B and C in invasion rather than co-culture. The red line and black line overlap at long times ($T_B = A_B$) because the ciliates eventually induce all bacteria to aggregate.

#### Predation rates and functional form

Here we justify three modeling assumptions regarding the predator-prey interaction between C and B: (1) the functional response (B loss term) is linear in bacterial densities, (2) the numerical response (growth of C due to predation on B) is negligible, (3) the feeding rate is estimated to be $1 \times 10^{-5}$ mL h$^{-1}$.

First, there is substantial evidence that the functional response of *T. thermophila* consuming bacterial prey is sigmoidal (e.g. $f_{max} x_B x_C/(U + x_B)$ where $f_{max}$ and $U$ are constants). The typical justification for this is the presence of a prey handling time which limits the absolute rate at which a predator can consume a prey [12]. Measurements of C feeding rates as a function of bacterial density show clear saturation at higher bacterial densities [13]. The sigmoidal dependence of ciliate uptake rates on prey concentration is further supported by studies of ciliate uptake of latex microspheres [14]. These studies give a half-velocity constant ($U$) for the sigmoidal prey uptake rate in the case of *T. thermophila* of $10^7$ bacteria mL$^{-1}$. Further studies show limited growth of *T. vorax* for *E. coli* densities below $2 \times 10^7$ mL$^{-1}$[15]. Therefore, the literature supports the conclusion that in our experiment, where bacterial densities never rise above $4 \times 10^6\,\mathrm{mL}^{-1}$, the functional response is well approximated by a linear model ($x_B << U$). We therefore use the term $F x_B x_C$ to describe the impact of C on the (planktonic) bacterial densities. In this linear model, following previous convention [14] we refer to the feeding rate as the per ciliate uptake rate of bacteria at low bacterial densities e.g. $F = f_{max}/U$.

Second, we claim that the feeding rate of ciliates on bacteria ($F$) has a value of $1 \times 10^{-5}$ mL h$^{-1}$. This claim is supported again by the literature. Fenchel estimated a feeding rate of *Tetrahymena* of approximately $10^{-5}$ mL h$^{-1}$ [9]. Hatzis *et al.* measured uptake rates in *Tetrahymena* of fluorescently labeled 2.74 µm diameter latex beads (at low bead concentration) and found rates between $10^{-5}$ to $10^{-4}$ mL h$^{-1}$ while noting substantial population level heterogeneity in bead uptake [10]. We note that studies with passive particles (latex spheres) avoid possible artifacts from prey aggregation. In a follow-up study the same authors note that the fraction of feeding ciliates declines substantially from about 80 % to 20 % as the ciliates enter stationary phase [11]. This is likely a contributor to the dynamics we observe in our study. However, we neglect this time dependent feeding rate in order to keep the model simple. Finally, we can make crude estimates of this rate from our data as well. If we neglect aggregation and algae, the dynamics of bacteria are given by $\dot{x}_B/x_B = (r_B - Fx_C)$. We measure $r_B$ =0.3 h$^{-1}$. We note that the growing bacteria are limited in their maximum density due to predation, at the crossover point (when predation and growth are balanced) $r_B = Fx_C$, or $F = r_B/x_C$. If we assume that this crossover point occurs when $x_C \sim 1 \times 10^4$ mL$^{-1}$ (and before bacteria have consumed all substrate) this gives an estimate of $F \approx 3 \times 10^{-5}$ mL h$^{-1}$, in good agreement with previous estimates. Due to the dependence of C feeding rates on growth state of the population and particle sizes, both of which are changing in our experiment as C enters stationary phase and B aggregates, we fixed the feeding rate on the lower end of the reported range: $10^{-5}$ mL h$^{-1}$. Note that this feeding rate only applies to planktonic bacteria, $x_B$. In order to replicate the inability of ciliates to eat aggregates of bacteria, there is no feeding of ciliates on aggregating bacteria $A_B$ in the model.

Third, we claim that the numerical response of the predator in response to predation is negligible and we therefore make C abundance dynamics independent of $x_B$ (Eqn. 4). The results of Seta and Tazaki [15] show no growth of *T. vorax* on *E. coli* when the abundance of the latter is below $2 \times 10^7$ mL$^{-1}$. In fact, these authors estimate a ciliate yield of $4 \times 10^4$ bacteria/ciliate (e.g. one ciliate is produced from the consumption of $4 \times 10^4$ bacterial cells). Further, Curds and Cockburn [13] measure the dry weight of *T. pyriformis* ($1.3 \times 10^{-10}$ g cell$^{-1}$ and the yield on bacteria (*Klebsiella aerogenes*) to be 50 % by dry weight. If we take the dry weight of *E. coli* to be 280 fg (`bionumbers.hms.harvard.edu`BNID: 103904) we estimate approximately $1 \times 10^3$ bacteria/ciliate. Therefore, we expect the yield of C on B to be between $1 \times 10^3$ to $4 \times 10^4$ bacteria/ciliate. Our data show that C predation reduces bacterial abundances from approximately $1 \times 10^6$ to $1 \times 10^5$ mL$^{-1}$ (Fig. S8c). Based on this reduction in B density, and the range of yields, we expect predation to produce between 50 and 1000 ciliates. Given the carrying capacity of C on this medium is $1.2 \times 10^4$ mL$^{-1}$ we conclude that the numerical response generates at most 10 % of the maximum C population. Finally, when we compare abundance dynamics of C in the presence and absence of B we see no significant difference (Fig. S21). These results support the modeling decision to omit any numerical response from our description of the community dynamics.

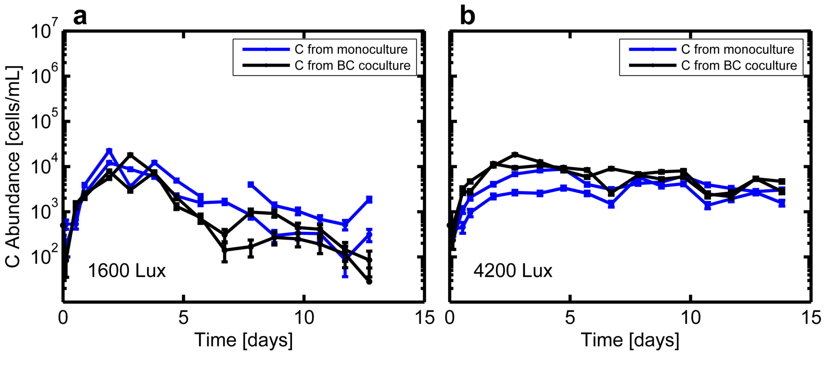


FIG. S21: Abundance dynamics of C in monoculture and in co-culture with B at 1600 Lux (**a**) and 4200 Lux (**b**).

#### Algae-bacteria interactions

Modeling AB interactions and dynamics requires three assumptions: (1) A growth rate depends only on light level and composition of the community, (2) A inhibits growth of B but not carrying capacity and (3) the de-aggregation rate of B due to the presence of A takes a value of $\alpha_2 = 2 \times 10^{-8}$ mL h$^{-1}$.

First, Fig. S1 shows measured growth rates of A as a function of light level and community composition. At low light algal growth rate decreases substantially with the addition of B and/or C. At high light, to within the precision of our measurement, A growth rate does not depend on community composition. Rather than construct a functional form with its own parameters that relates algal growth rate to light and community composition, we simply set the algal growth rate for each combination of light level and community composition. Since the dynamics of B are the focus of our study, and A dynamics are always well described by a logistic model, this modeling choice is well justified and removes unnecessary parameters from our model.

Second, as shown in Eqn. 1, we assume that A impacts the growth rate of B in a density dependent fashion ($r_B S - r_{AB} x_A / K_A$), but also that A has no impact on the carrying capacity of B. Our data support this assumption since we see no impact of the presence of A on the carrying capacity of B. In the situation where B invades a high density ($>1 \times 10^5$ mL$^{-1}$) algae culture, we find that the bacterial growth is strongly inhibited in about half of the cases we observed. We therefore capture this growth inhibition using the term shown above. We note that this purely deterministic model will not capture the stochastic outcomes we observe experimentally (that would require a more detailed model of the A inhibition of B and likely the predation process). We set $r_{AB}$ to be on the same order as $r_B$. We make this assumption since our data clearly show that A is capable of nearly completely, or completely, inhibiting the growth of B (e.g. when $x_A = K_A$ near-complete inhibition will occur if $r_{AB} \approx r_B$).

Third, we must specify the unknown parameter $\alpha_2$. To do so, we treat it as a free parameter in our simulation since we have no direct measure of the rate at which A induces B to de-aggregate. We performed three simulations of the full three species (ABC) community: AC, high light $t_{inv} = 4$ d; ABC high light, co-culture; AC low-light,$t_{inv} = 4$ d. For each condition we swept $\alpha_2$ over a range of logarithmically spaced values. Our expectation is that for a single value of $\alpha_2$ we should find that B fails to invade in AC high light at 4 d, successfully grows in co-culture with AC, and successfully invades AC low light at 4 d. The results of these three simulations are shown in Figs. S22,S23,S24. From these simulations we find that $\alpha_2$ =$2 \times 10^{-8}$ mL h$^{-1}$ captures the experimentally observed dynamics in all three conditions (see panels with red axes in Figs. S22,S23,S24).

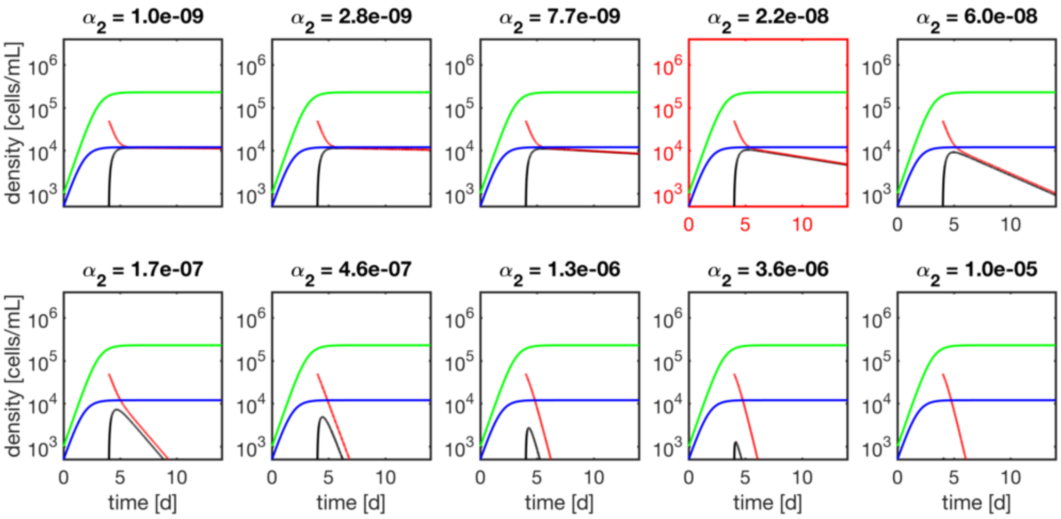


FIG. S22: Abundance dynamics of A, B and C in high light conditions with $t_{inv} = 4$ d. Parameter values are as shown in Table I with the exception of $\alpha_2$ which is varied as indicated in the title of each panel. In each panel $x_A$ is in green, $x_C$ in blue, $T_B$ in red, and $A_B$ in black. The units of $\alpha_2$ are mL h$^{-1}$.

##### Aspects of the dynamics not captured by the model

The model presented here is necessarily simplified to limit the number of free parameters in the simulation to one ($\alpha_2$). As a result there are several qualitative aspects of the data that are not captured by the modeling framework. Here we enumerate these.

- B exhibits a transient spike in aggregation as it enters stationary phase even in the absence of A or C. (e.g. Fig. S3g). This process has been examined previously in our group and explained using a substrate dependent aggregation process.[16] We neglect this aspect of the bacterial aggregation dynamics.

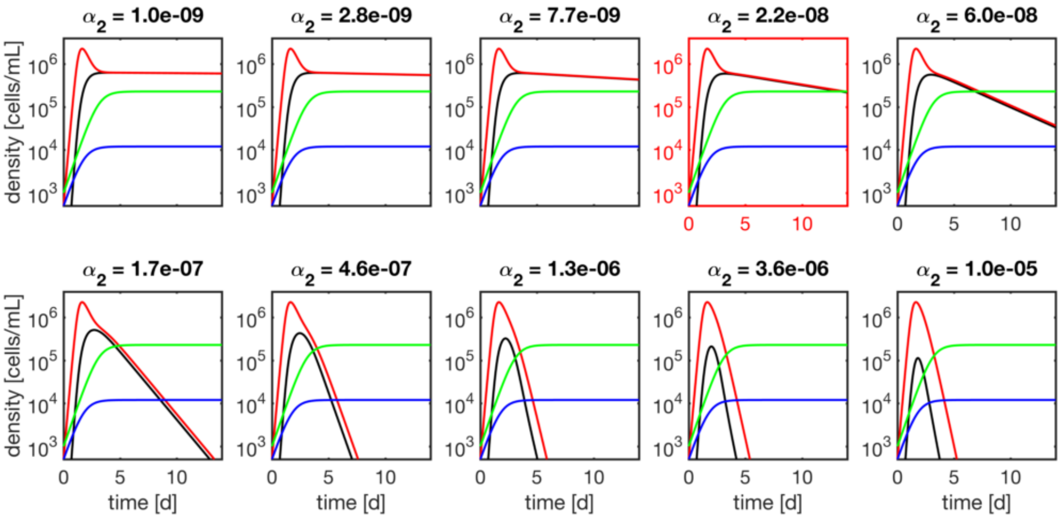


FIG. S23: Abundance dynamics of A, B and C in high-light, co-culture conditions simulated using the model discussed above. Parameter values are as shown in Table I with the exception of $\alpha_2$ which is varied as indicated in the title of each panel. In each panel $x_A$ is in green, $x_C$ in blue, $T_B$ in red, and $A_B$ in black. The units of $\alpha_2$ are mL h$^{-1}$

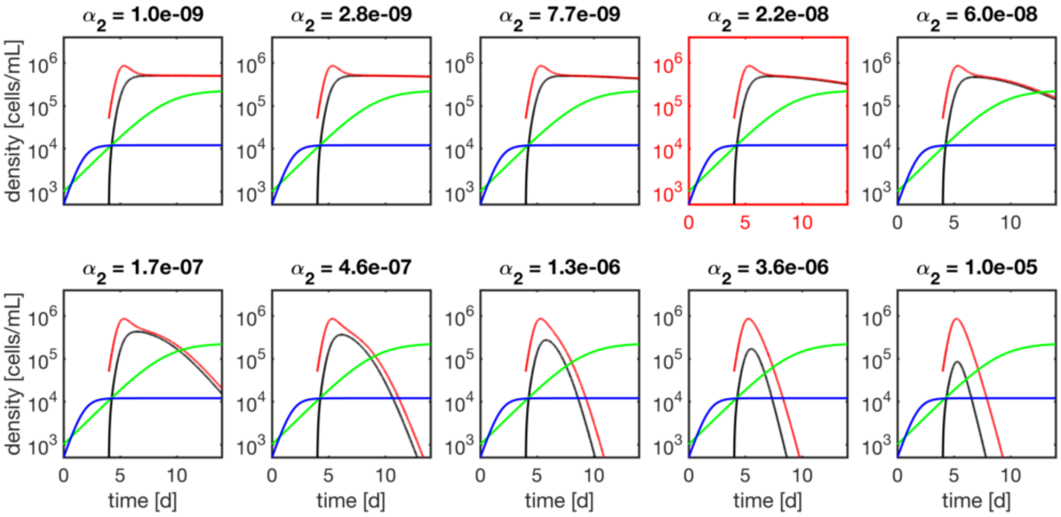


FIG. S24: Abundance dynamics of A, B and C in low light conditions for $t_{inv} = 4\,\mathrm{d}$ simulated using the model discussed above. Parameter values are as shown in Table I with the exception of $\alpha_2$ which is varied as indicated in the title of each panel. The growth rate for A in this simulation is taken to be the low light growth rate in the presence of C alone ($r_A$ =0.025 h$^{-1}$). In each panel $x_A$ is in green, $x_C$ in blue, $T_B$ in red, and $A_B$ in black. The units of $\alpha_2$ are mL h$^{-1}$

- In co-culture conditions with B in the presence of C, after the peak in bacterial abundance and then subsequent fall due to predation, the abundances of B rise over the last ∼10 days of the experiment. (Fig. 6 h, main text). This rise could be explained by slow growth of aggregated bacteria or by growth of B on the detritus of dying C. Our model fails to capture the rise and we have not included it since we have no direct evidence for either of these processes.

- The deterministic ODE framework does not capture the stochasticity we observe in the outcome of B invading A alone (Fig. S9). This stochasticity could be captured by an effective randomness in the parameter $r_{AB}$. We have not included it here since the process is likely driven by population structure in either A or B which is not present in the current model (e.g. phenotypic heterogeneity in the response of B to inhibition by A). In the absence of direct mechanistic insight into this stochasticity process, we omitted it from our model.

- The decline in C abundances after approximately 4 days is not modeled here. This decline could be addressed by inferring a death rate from the data, but it would not qualitatively impact the agreement between the model and the simulation.
